# A human fetal lung cell atlas uncovers proximal-distal gradients of differentiation and key regulators of epithelial fates

**DOI:** 10.1101/2022.01.11.474933

**Authors:** Peng He, Kyungtae Lim, Dawei Sun, Jan Patrick Pett, Quitz Jeng, Krzysztof Polanski, Ziqi Dong, Liam Bolt, Laura Richardson, Lira Mamanova, Monika Dabrowska, Anna Wilbrey-Clark, Elo Madissoon, Zewen Kelvin Tuong, Emma Dann, Chenqu Suo, Isaac Goh, Masahiro Yoshida, Marko Z Nikolić, Sam M Janes, Xiaoling He, Roger A Barker, Sarah A Teichmann, John C. Marioni, Kerstin B Meyer, Emma L Rawlins

## Abstract

We present a multiomic cell atlas of human lung development that combines single cell RNA and ATAC sequencing, high throughput spatial transcriptomics and single cell imaging. Coupling single cell methods with spatial analysis has allowed a comprehensive cellular survey of the epithelial, mesenchymal, endothelial and erythrocyte/leukocyte compartments from 5-22 post conception weeks. We identify new cell states in all compartments. These include developmental-specific secretory progenitors and a new subtype of neuroendocrine cell related to human small cell lung cancer. Our datasets are available through our web interface (https://lungcellatlas.org). Finally, to illustrate its general utility, we use our cell atlas to generate predictions about cell-cell signalling and transcription factor hierarchies which we test using organoid models.

**Highlights:**

- Spatiotemporal atlas of human lung development from 5-22 post conception weeks identifies 144 cell types/states.
- Tracking the developmental origins of multiple cell compartments, including new progenitor states.
- Functional diversity of fibroblasts in distinct anatomical signalling niches.
- Resource applied to interrogate and experimentally test the transcription factor code controlling neuroendocrine cell heterogeneity and the origins of small cell lung cancer.

## Introduction

Single cell mapping of cell states in the adult human lung in health and disease is being performed at increasing resolution (Carraro and Stripp 2022) providing a foundation for understanding lung cellular physiology. The adult lung has low rates of cell turnover (Blenkinsopp 1967; Rawlins and Hogan 2008) making it difficult to capture transition states and progenitor cells. Moreover, there are developmental-specific cell states that do not exist in the adult. A high-resolution cell atlas of the embryonic and fetal human lung will identify developmental precursors and progenitors, predict differentiation trajectories and potential gene regulatory networks. This will provide a baseline for studying adult homeostasis and disease.

The lung buds are specified in the human foregut endoderm at ∼5 post conception weeks (pcw) (Burri 1984; Nikolić, Sun, and Rawlins 2018). Subsequent morphogenesis is driven by branching of the distal-most bud tips. The bud tip epithelium comprises SOX9^+^, ID2^+^ multipotent progenitors which self-renew during branching (Rawlins, Clark, et al. 2009; Alanis et al. 2014; Nikolić et al. 2017; Miller et al. 2018). As the bud tip epithelium branches into the surrounding mesoderm, the epithelial cells that remain in the stalk region start to differentiate into bronchiolar (airway) epithelium (∼5-16 pcw) and then (from ∼16 pcw) into alveolar epithelium (Nikolić, Sun, and Rawlins 2018). The pattern of growth from undifferentiated multipotent epithelial progenitors at the distal tips means that the position of a cell along the proximal-distal axis of the lung epithelial tree is a strong predictor of its maturity. The more mature cells, which exited the tip first, are found more proximally, whereas the most immature cell states, which exited the tip recently, are found in the tip-adjacent (stalk) regions (Rawlins et al. 2007). In other words, space reflects time in lung development. Therefore, coupling single cell state identification to *in vivo* spatial visualisation can provide high confidence in the identification of novel progenitor cells in the developing lung. Moreover, detailed spatial analysis of cell states allows cell identity designations to be compared to more traditional histological definitions.

We have generated a high-resolution single cell atlas of human lung development using a combination of scRNA-seq, scATAC-seq, Visium Spatial Transcriptomics, and mRNA *in situ* hybridisation using hybridisation chain reaction (HCR) (Trivedi et al. 2018). Combining these data sources has allowed us to identify 144 cell states/types in 5-22 pcw lung samples. These include novel progenitor cell states, transition populations and a new subtype of neuroendocrine cell which is related to a subtype of human small cell lung cancer. We observe increasing cell maturation over time with many cell states identified in the adult human lungs already present at 22 pcw. Moreover, we have used our atlas to make predictions about progenitor cell states, signalling interactions and lineage-defining transcription factors, and we demonstrate how these can be efficiently tested using a genetically-tractable human fetal lung organoid model. These data sets are available for interactive analysis at https://lungcellatlas.org.

## Results

### A single cell atlas of human lung development comprising 144 cell states

We obtained human embryonic and fetal lungs from 5-22 pcw for scRNA-seq and scATAC-seq. To focus on differentiation we deeply sampled 15, 18, 20 and 22 pcw lungs and separated proximal and distal regions, while leaving lungs at 5, 6, 9 and 11 pcw intact. We used a mixture of cell dissociation methods to obtain a balanced mixture of cell types (Fig. 1A) and produced high-quality transcriptome (Fig. S1A; average >2400 genes/cell) and DNA accessibility (Fig. S1O,P; average >18,000 fragments/nucleus) data. The RNA profiles of cells from different dissociation protocols and compartments were iteratively parsed out by clustering (Fig. S1C) and subclustering (Fig. S1D) without batch correction to maintain biological features. To enrich biological features and mitigate technical ones, we removed doublet-driven clusters (Fig. S1E,G,H,K), stressed or low-quality clusters (except those expressing known markers, such as erythroid) (Fig. S1I,J,N), clusters composed of cells from only one sample when replicates are available, and clusters of cells from other organs due to contamination during dissection (Fig. S1L) (He et al. 2020; Cao et al. 2020). We identified maternal cells, which are a minimal fraction in our atlas (Fig. S1M), based on genetic background (Fig. S1F). The result of this clustering is shown as a Uniform Manifold Approximation and Projection (UMAP) (Fig. 1A), on which we manually annotated fibroblast, epithelial, endothelial and erythrocyte/leukocyte lineages (Fig. 1B). Plotting the cell type distribution against time (excluding trypsin/CD326-treated samples, shown in Fig. S1B), showed that fibroblasts were the most prominent cell, particularly in younger lungs (Fig. 1C). Leukocytes and erythrocytes were observed in all lungs sampled, with B, T and NK cells becoming prominent from 15 pcw (Fig. 1C).

**Figure 1.**
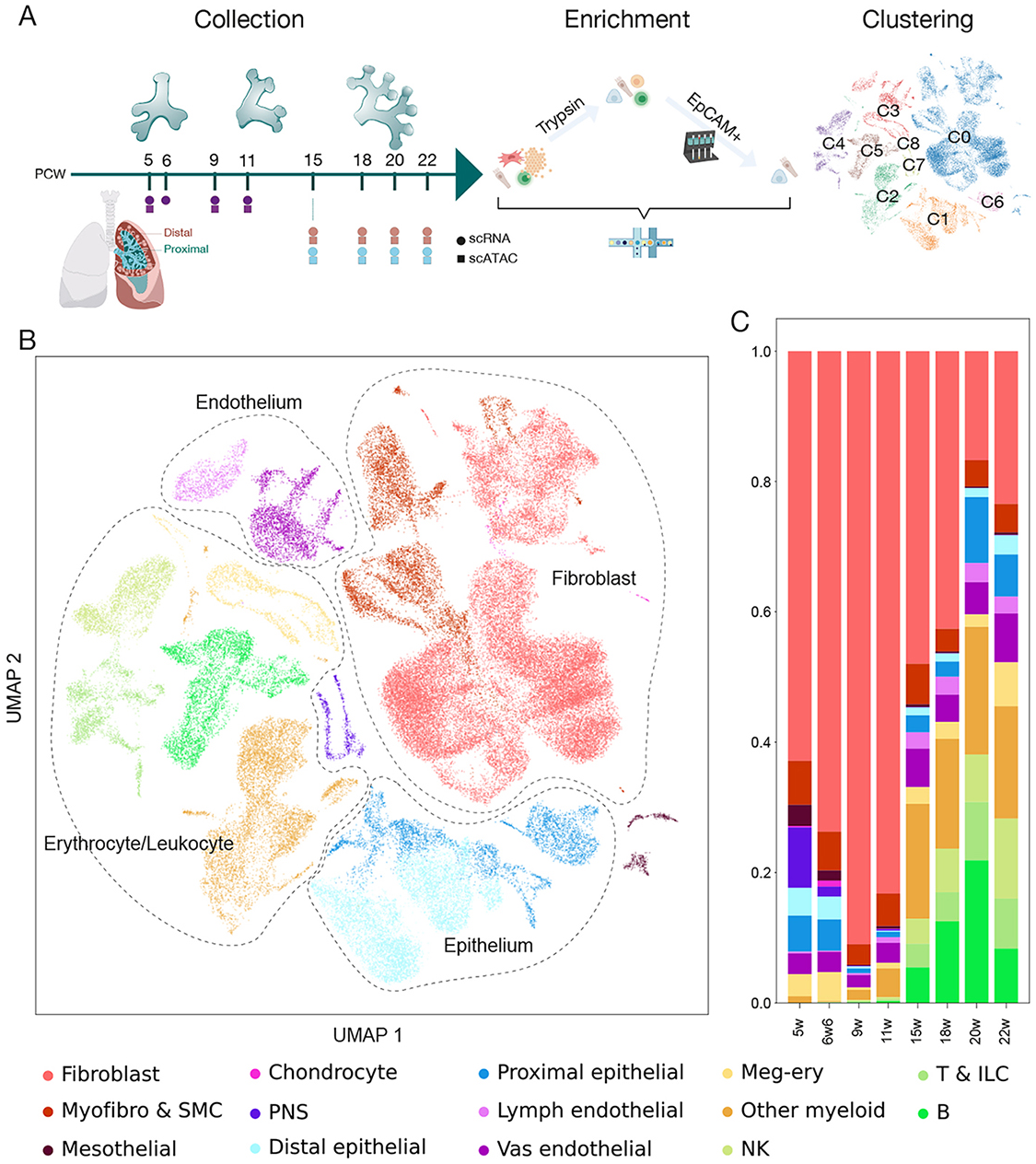
Experimental overview. (A) Overview of sample collection for scRNA-seq (circles) and scATAC-seq (squares) experiments from whole lung (purple), distal (red) and proximal (blue) regions, cell processing and broad clustering; cluster number refers to the data portal (https://lungcellatlas.org). (B) UMAP representation of ∼80,000 good quality cells, indicating epithelial, endothelial, fibroblasts and leukocyte/erythroid compartments. (C) Cell type proportions of the whole lung over developmental time. See also Figures S1, S2, S3 and S9.

Further cell type annotation was performed via sub-clustering (see methods) and cluster naming was based on observed and published marker genes (Supplemental Table 1), resulting in assignment of 144 cell types/states (Fig. S2A). Sample age was a strong determinant of clustering (**χ**^2^=163727, p ≈ 0), reflecting progressive cell type maturity over time (Fig. S2B). Clusters mostly grouped into three distinct regions which we categorised as early (5, 6 pcw), mid (9, 11 pcw) and late (15-22 pcw) stages. Cell cycle phase (Fig. S2C, **χ**^2^=25361, p ≈ 0) and dissected region (Fig. S2D, **χ**^2^=968, p = 8.9E-131) were also associated with clustering. However, region was only prominent for a small number of proximally-located cell types (Fig. S2H), suggesting that most proximal to distal regions of the airway branch structure were still represented in both dissected regions of the lung. Epithelial cells were mostly derived from the trypsin-treated and CD326-enriched samples, although airway smooth muscle, myofibroblasts and alveolar fibroblasts were also enriched here (Fig. S2E). Peripheral Nervous System (PNS) cells and chondrocytes were only obtained from 5-6 pcw lungs, likely correlating with lower extracellular matrix (ECM) complexity and/or increased fragility of older neurons. PNS cells were clustered and assigned to cell types, but their scarcity precluded further analysis (Fig. S2A,F,G). Data integration and logistic regression-based comparison showed that gene expression of our annotated cells corresponds well to those from adult lungs (Madissoon et al. 2021) (Fig. S3A-C).

### A differentiation trajectory of airway progenitor states lies along the distal to proximal axis of the developing lungs

The epithelial cells separate by age (Fig. 2A,B), with many basal cells, MUC16^+^ ciliated cells and secretory cells enriched in the proximally-dissected tissue (Fig. 2B; S2H). The most immature epithelial progenitors are tip cells: SOX9^+^ multipotent progenitors located at the distal branching tips of the respiratory tree (Nikolić et al. 2017). Tip cells were separated by developmental age into early (5,6 pcw), mid (9,11 pcw) and late (15-22 pcw) populations (Fig. 2A,B) with some shared and some stage specific markers (Fig. 2C). On the epithelial UMAP, each tip population clusters closely with adjacent stalk cells (*SOX9^LO/-,^ PDPN^LO^, HOPX^LO^*) and airway progenitors (*CYTL1^LO/+^, PCP4^+^, SCGB3A^+/LO^*) (Fig. 2A). The tip, stalk and airway progenitors can be visualised in a distal-proximal sequence in the tissue at all stages tested (10-16 pcw) (Fig. 2E; S4A,B, Supplemental video 1), consistent with the most proximal cells being the most mature. These three cell types form a predicted differentiation trajectory from mid-tip to mid-stalk to mid-airway progenitor which branches into the neuroendocrine, or secretory, lineages (Fig. 2D).

**Figure 2.**
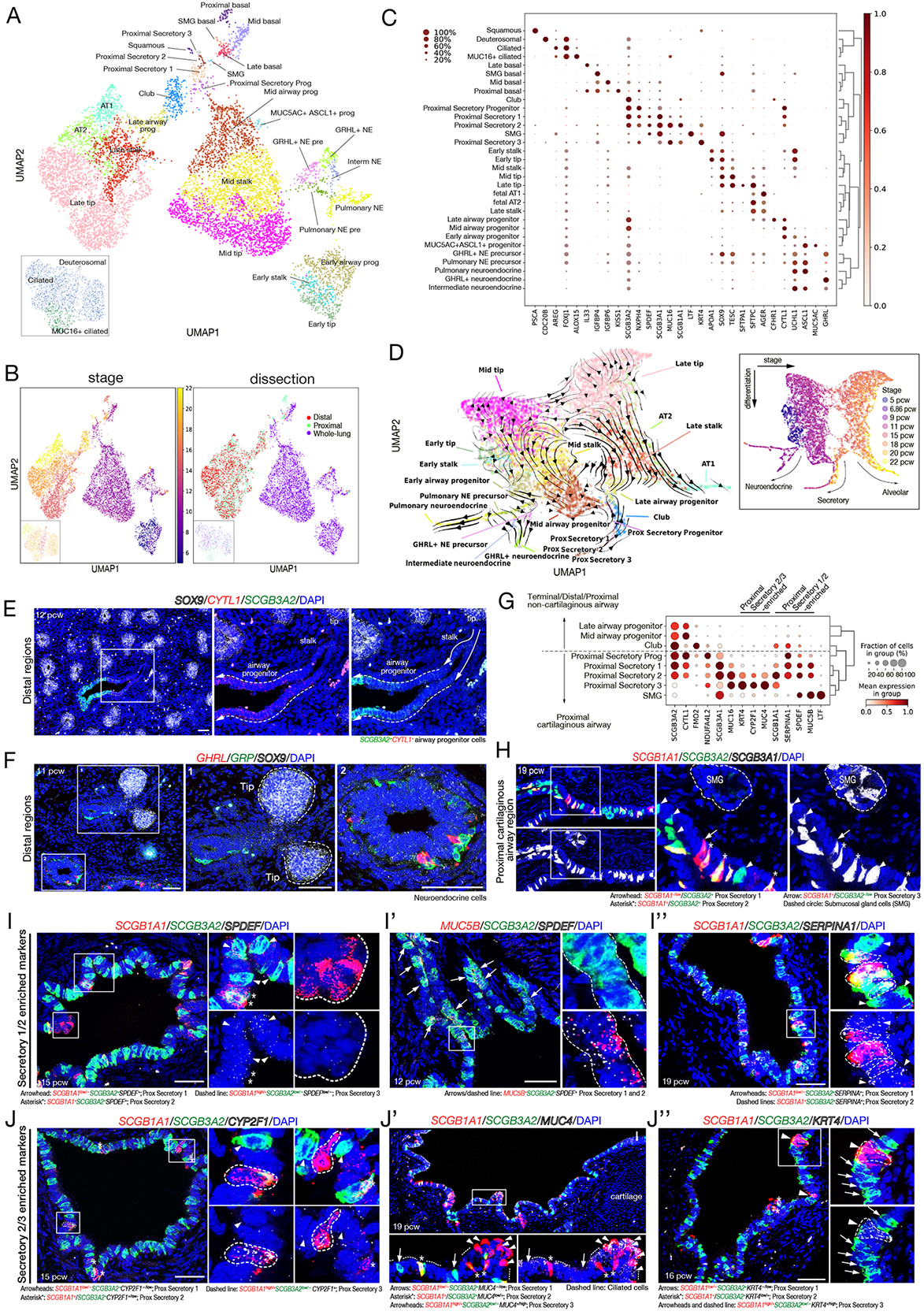
Epithelial cell types, states and location over developmental time. (A, B) UMAP visualisation of epithelial cells, coloured by cell types (A), stage (B, *left*), and region (B, *right*). (C) Dot plot describing differential marker gene expression level for epithelial cells. (D) UMAP visualising the predicted epithelial cell lineage trajectory using scvelo, inset: developmental age. (E, F) *In situ* HCR at 11 (F) and 12 (E) pcw. (E) *SOX9* (tip epithelium, white), *CYTL1* (red), *SCGB3A2* (green). (F) *GHRL*^+^ (*GHRL*^+^ neuroendocrine, red), *GRP*+ (pulmonary neuroendocrine, green). (G) Dot plot showing differential marker genes across secretory cell subtypes. (H) *In situ* HCR at 19 pcw using *SCGB1A1* (red), *SCGB3A2* (green), and *SCGB3A1* (white). (I-J) Differentially enriched genes in the proximal secretory cell subtypes. *SPDEF* (I, I’), *MUC5B* (I’), and *SERPINA1* (I’’), *CYP2F1* (J), *MUC4* (J’), *KRT4* (J’’) all white; *SCGB1A1* (red) and *SCGB3A2* (green). DAPI, nuclei. Scale bars, 50 μm. See also Figures S4, S5 and S6.

### Two subtypes of neuroendocrine cells are present in the developing airways

Consistent with previous data (Cutz, Gillan, and Bryan 1985), the earliest differentiated epithelial cells detected were neuroendocrine (NE) cells in 5 pcw lungs (Fig. 2A-C). We identified two types of NE cells: classical pulmonary NE cells (*GRP^+^*) and GHRL^+^ NE cells (*TTR^+^*, *GHRL^+^*) in agreement with a recent human fetal cell atlas (Cao et al. 2020). We observed increasing maturity of NE cells over time (specific populations denoted as precursors on the UMAP). In addition, an intermediate NE population, a putative transition state, connected the two NE cell types (Fig. 2A). At 11 pcw, *GRP^+^* NE cells were always observed closer to the budding tips, suggesting that they begin to differentiate prior to the *GHRL^+^*NE cells (Fig. 2F). This spatial difference was not apparent in the oldest samples where both *GRP^+^*and *GHRL^+^* cells were observed at all airway levels, although less abundant distally (Fig. S4C). Mouse GHRL^+^ NE cells were not detected in re-analysis of published mouse data (Negretti, Plosa, Benjamin, Schuler, Christian Habermann, et al. 2021; Zepp et al. 2021), or spatially (Borromeo et al. 2016). However, *Ghrl* is expressed in mouse ciliated cells which cluster with human fetal GHRL^+^ NE cells following scVI integration (Lopez et al. 2018) and clustering analysis (Fig. S3D-F) (Zepp et al. 2021).

### Multiple secretory cell subtypes in the proximal cartilaginous airways

We annotated 5 sub-types of differentiating secretory cells and one more immature proximal secretory progenitor. (i) The proximal secretory progenitors (*SCGB3A2^+^, SCGB1A1^-^, SCGB3A1^-^_/LO_, CYTL1^+^*) were detected in the single cell atlas at 9 pcw, prominent at 11 pcw, but rarer in older lungs consistent with a progenitor state (Fig. 2A-C,G). (ii) Club cells (S*CGB3A2^+^, SCGB1A1^+^, SCGB3A1^-^, SPDEF^-^, MUC16^-^*) were detected from 15 pcw in the single cell data (Fig 2A-C,G), or 12 pcw in the tissue localised in clusters more distally, but largely dispersed in the more proximal non-cartilaginous regions (Fig. S4D). (iii) Submucosal gland (SMG) secretory cells (*LTF^+^, SCGB3A1^+^, SPDEF^+^*) were detected from 15 pcw in the single cell data, located in SMG ducts and likely to be a precursor of serous and/or mucous-secreting SMG cells (Fig. 2A-C,G; S4G). (iv) Proximal secretory 1 (*SCGB1A1^LO^, SCGB3A2^+^, SCGB3A1^+^*) and (v) proximal secretory 2 (*SCGB1A1^+^, SCGB3A2^+^, SCGB3A1^+^*) appeared from 11 pcw (Fig. 2A-C,G-H; S4E). Both were *SPDEF^+^*, *MUC5B^+^, SERPINA1^+^*(Fig. 2I), suggesting they differentiate into goblet or mucous cells. By contrast, (vi) proximal secretory 3 (*SCGB1A1^+^, SCGB3A2^LO/-^, SCGB3A1^+^*) was detected from 15 pcw and was *SPDEF^-^* (Fig. 2A-C), but *CYP2F1^+^, MUC4^+^* and *KRT4^+^* (Fig. 2G,J). All three luminal proximal secretory cell populations were located in the proximal cartilaginous airways and were *MUC16^+^* (Fig. 2C,G,H; S4E,F). Detailed spatial-temporal analysis of 10-21 pcw airways revealed that the proportion of proximal secretory progenitors decreased with developmental age, whilst proximal secretory cells 1 and 2 increased (Fig. S5A-C); consistent with a progenitor function for proximal secretory progenitors.

### Other airway cells

We detected ciliated cells (*FOXJ1^+^, ALOX15*^+^) from 11 pcw, interspersed with secretory/club cells throughout the airways (Fig. 2A-C; S4H; S5A,B). Rarer deuterosomal cells (*FOXJ1^+^, CDC20B^+^*) appeared at the same time (Fig. 2A-C). MUC16^+^ ciliated cells (*FOXJ1^+^*, *DNAH^+^, MUC16^LO^*) were also detected from 11 pcw, but confined to proximal dissected regions (Fig. 2A-C; S2H). They were located in patches in the most proximal cartilaginous airways (Fig. S4I), and likely represent *MUC16^+^* secretory cells generating ciliated cells, as suggested in the adult (Deprez et al. 2020; Carraro et al. 2021; Vieira Braga et al. 2019). Basal cells (*TP63^+^, F3^+^*) were present from 9 pcw (Fig. 2A-C; S4J) and more frequent in proximal regions (Fig. 2C; S2H; S5A,B). Rarer cells (ionocytes, tuft) that have been consistently identified in adult airways were not present in our single cell data. However, we found putative ionocytes (*FOXI1^+^*; 4/4 lungs) and putative tuft cells (*POU2F3^+^*; 2/4 lungs) in the most proximal cartilaginous airways of 21-22 pcw lung sections (Fig. S5E), suggesting they begin to differentiate mid-gestation. Moreover, we reproducibly detected a small population of *MUC5AC*^+^, *ASCL1*^+^ cells in 9,11 pcw lungs (Fig. 2A-C). These were localised to the proximal non-cartilaginous airways where they appeared as solitary, somewhat basal, non-columnar cells (Fig. S4K). We hypothesise that they are an unknown progenitor, consistent with their transient appearance and the observation that Ascl1^+^ NE cells in adult mice can generate club, ciliated and mucous cells following injury (Yao et al. 2018; Ouadah et al. 2019).

### Predicted airway epithelial differentiation trajectories

A detailed spatio-temporal analysis of the major airway epithelial cell types from 10-21 pcw confirms that cell maturation begins in the more proximal regions. An example is lack of ciliated and club cells in the distal non-cartilaginous airways at 10-12 pcw, but presence at 15-21 pcw (Fig. S5A-D). Conversely, airway progenitors are found throughout the non-cartilaginous airways at 10-12 pcw, but restricted to the terminal airways by 15-21 pcw (Fig. S5A-D). In addition, proximal secretory cells are spatially restricted to the cartilaginous airways, whilst club cells are found in the non-cartilaginous regions (Fig. S5A-D).

This spatial separation means that predicted differentiation trajectories which combine proximal secretory cells and club cells (Fig. 2D) can reveal general trends, but are likely to be over-simplified. We therefore predicted mid- (Fig. S6A-C) and late-stage (Fig. S6D-F) airway lineage trajectories separately. In both cases, basal cells formed discrete clusters on the UMAPs (Fig. S6A’,D’). Trajectory inference analysis suggests a differentiation route from mid-tip to stalk to airway progenitors to proximal secretory progenitors and proximal secretory cells (Fig. S6B), consistent with sample age (Fig. S6B’). Visualising gene expression along the inferred trajectory shows mid-tip and stalk cells are similar (Fig. S6C). The stalk cells lose some tip markers, including *FOXP2* and *SOX9*, and gain a relatively small number of genes including *PDPN* and *AGER*. By contrast, the newly-defined airway progenitors upregulate marker genes associated with airway fates, including *CYTL1*, *CLDN4* and *SCGB3A2* (Kaarteenaho et al. 2010; Guha et al. 2012) (Fig. S6C). A similar differentiation trajectory was predicted from late-tip to late-stalk to late-airway progenitor to club cells (Fig. S6E), although the oldest tip and stalk cells included in this analysis may produce alveolar lineages (Fig. S6E’; 3C-E; S7A,B). Visualising gene expression along the inferred late-airway trajectory shows that the late-tip and stalk cells are transcriptionally similar and undergo gene expression changes analogous to mid-tip and stalk (loss of *SOX9*, *FOXP2*; gain of *PDPN*, *AQP5*; Fig. S6F).

These analyses predict that cells first exit the tip into a tip-adjacent stalk-state, followed by gain of airway progenitor identity before commitment to a specific differentiation state that likely depends on local signalling cues. Although we cannot predict the origin of the basal cells using trajectory inference methods, we hypothesise that they are derived from a columnar progenitor (possibly the airway progenitor), but will themselves act as progenitor/stem cells following differentiation analogous to previous observations in mice (Yang et al. 2018).

Our trajectory inference (Fig. S6A-F) predicts that airway progenitors will differentiate readily to airway cell types. At 9-10 pcw, *CYLT1^+^*, *SCGB3A2^+^* airway progenitors are found throughout the airway tree (Fig. 2E; S4B,D,L; S5A,B,D). We devised a strategy to isolate airway progenitors using a combination of distal non-cartilaginous airway micro-dissection and transduction with a lentiviral *SCGB3A2* transcriptional reporter (*SCGB3A2-GFP*, Fig. S6G). Freshly-isolated distal *SCGB3A2*-GFP^+^ cells were *SOX9^LO^*, *CYTL1^HI^*, *SCGB1A1^LO^*, *SCGB3A2^LO^*, and *SCGB3A1^LO^* compared to tip/stalk cells and more proximal *SCGB3A2*-GFP^+^ cells from the same lungs (Fig. S6H), confirming airway progenitor identity. When single cells were placed into an FGF-containing differentiation medium (Hawkins et al. 2021), distal *SCGB3A2*-GFP^+^ cells produced basal, ciliated and mature secretory cells (Fig. S6I-K). This demonstrates that, consistent with the trajectory analysis, the airway progenitors are competent to differentiate into airway lineages.

In summary, we have identified multiple epithelial progenitor states (tip; stalk; airway progenitor; proximal secretory progenitor) and differentiating airway cells which localise to a spatial differentiation gradient along the proximal-distal axis of the epithelium (summarised in Fig. S4L; S5D). Moreover, we identify *GHRL^+^*neuroendocrine cells which do not exist in the mouse.

### Late epithelial tip cells acquire alveolar identity prior to alveolar epithelial differentiation

The tip cells expressed a core set of tip-specific markers (*SOX9^+^, ETV5^+^, TESC^+^, TPPP3^+^, STC1^+^*) at all stages sampled (Fig. 2A-C). We observed a gradual decrease in tip marker expression and an increase in alveolar type 2 (AT2) cell gene expression in tip cells with developmental age (Fig. 2C). Indeed, by 15 pcw the AT2 markers *SFTPC* and *SFTPA* were detected readily in the late-tip cells where they were co-expressed with lower levels of core tip markers (Fig. 3A,B). This late-tip is a unique tip cell transcriptional state which has not been detected in developing mouse lungs (Negretti, Plosa, Benjamin, Schuler, Habermann, et al. 2021; Zepp et al. 2021). The change in tip gene expression correlates with predicted differentiation trajectories from early to mid to late-tip cells to (late-stalk to) fetal AT2 and AT1 cell fates (Fig. 3C,D; without late-stalk in S7A). However, trajectory inference analysis at this transitional developmental stage is challenging. It is likely that some of the late-tip cells are producing the terminal branches of the conducting airways (Fig. S6D-F). Moreover, the inferred connections between mid-tip and late-tip cells are weak (Fig. 3C) and we cannot exclude a novel origin for late-tip cells perhaps emerging as new buds from a stalk position, although this hypothesis is not strongly supported by our analysis (Fig. 2D). Nevertheless throughout this period, similar to earlier stages, the late-tip cells remain SOX9^+^ and the late-stalk cells turn off tip markers and acquire *PDPN/AGER* (Fig. S4A; S7D).

**Figure 3.**
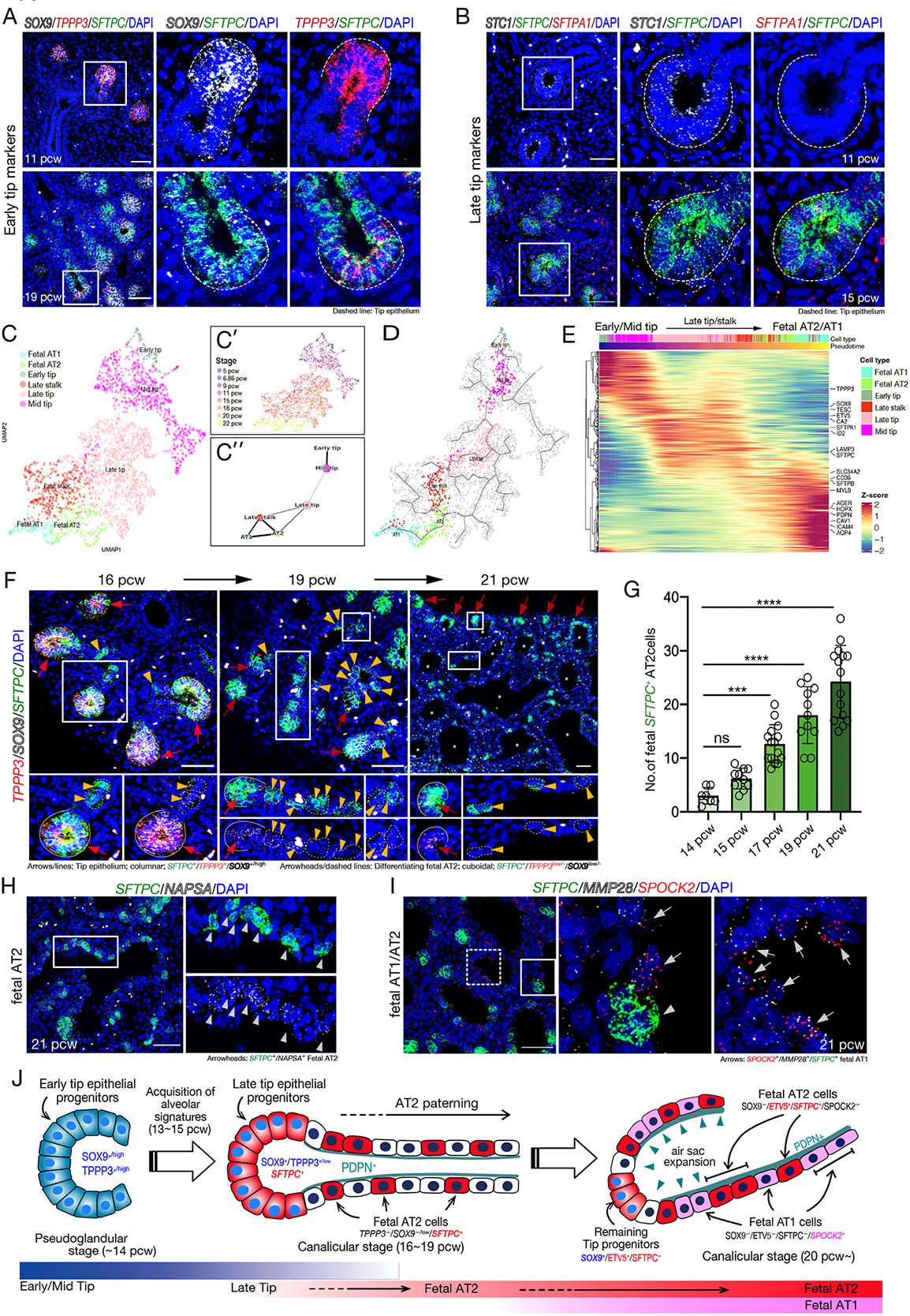
Late epithelial tip cells acquire an alveolar progenitor identity. (A, B) *In situ* HCR at 11 (A, B), 15 (B), and 19 (A) pcw. (A) *SFTPC* (green), *TPPP3* (red), *SOX9* (white). (B) *SFTPC* (green), *SFTPA1* (red), *STC1* (white). Dashed lines: tip epithelium. (C,D) UMAP visualisation of early to late tip, late stalk, fetal AT1 and AT2 cells, coloured by cell types (C) and stages (C’); PAGA analysis (C’’); Monocle3 trajectories (D). (E) Gene expression heatmap of trajectory coloured in D. (F) *In situ* HCR at 16, 19, and 21 pcw, *SFTPC* (green), *TPPP3* (red), and *SOX9* (white). White lines/red arrows: columnar tip prognitors, *SFTPC*^+^/*SOX9*^+/high^/*TPPP3*^+^. Arrowheads/dashed lines in stalk/air sac regions: cuboidal differentiating fetal AT2 cells, *SFTPC*^+^/*SOX9*^low/-^/*TPPP3*^low/-^. Asterisk (*) primitive air sacs. (G) Quantification of cuboidal *SFTPC*^+^/*SOX9*^low/-^ fetal AT2 cells in stalk/air sac regions in F. The *SFTPC*^+^ tip epithelial cells were excluded by their columnar morphology and marker expression (*SOX9*^low/-^). Mean ± SD, n >7. Significance evaluated by 1-way ANOVA with Tukey multiple comparison post-test; ns: not significant, \**P*<0.05, \*\**P*<0.01, \*\*\**P*<0.001, \*\*\*\**P*<0.0001. (H, I) *In situ* HCR analysis at 21 pcw. Fetal AT2 *SFTPC*^+^ and *NAPSA*^+^ (*arrowheads;* H, I) and fetal AT1 *SFTPC*^-^/*MMP28*^+^/*SPOCK2*^+^ (*arrow*s; I). (J) Diagram of the acquisition of alveolar progenitor identity by late epithelial tips, followed by differentiation to fetal AT2 and AT1 lineages. DAPI, nuclei. Scale bars, 50 μm. See also Figure S7 and S8.

A small number of AT2 cells appear in the single cell data from 15 pcw, but are more prominent from 22 pcw (Fig. 2A). Similarly, at 16 pcw late tip cells (*SOX9^+^,TPPP3^+^,SFTPC^+^*) were clearly visualised in the tissue, but differentiating AT2 cells (*SOX9^LO/-^,TPPP3^LO/-^,SFTPC^+^*) were rare suggesting AT2 production is just beginning (Fig. 3F,G; S7C). Over the following weeks, the size of the tip regions decreased and more differentiating AT2 cells (*SOX9^LO/-^,TPPP3^LO/-^,SFTPC^+^*) were detected (Fig. 3F,G). At 21 pcw smaller numbers of late tip cells persist and AT2 cells (*SOX9^-^, SFTPC^+^, NASPA^+^, ETV5^+^)* were found scattered throughout the developing air sacs (Fig. 3H, S7E-J). Consistent with the predicted change in tip fate potential (Fig. 3C-E), late-tip cells (16-20 pcw) grown as organoids retained a late-tip phenotype *in vitro* and were much more readily differentiated to mature AT2s than organoids derived from earlier developmental stages (Lim et al. 2021).

In our single cell atlas, differentiating AT1 cells were first visible at 18 pcw, but more prominent by 22 pcw (Fig. 2A-C). Similarly in tissue sections, AT1 cells were not detected at 17 pcw (Fig. S7H). However, by 20 pcw differentiating AT1 cells (*SPOCK2^LO^, SFTPC^-^*) were visible and at 21 pcw AT1 cells (*SPOCK2^+^, SFTPC^-^*) were interspersed with AT2 cells lining the developing air sacs (Fig. 3I; S7I,J). In sections, AT1 markers were only detected in cells which had undetectable, or extremely low levels of, *SFTPC* (Fig. S7H-J). Moreover, *SFTPC*-negative cells were always observed in the stalk regions from 16 pcw onwards (Fig. 3F). These spatial expression data are consistent with an alveolar epithelial differentiation model in which from ∼16 pcw the late-tip progenitors first exit the tip state, turning off co-expressed AT2 cell markers, and enter the late-stalk cell state, prior to initiating AT1 or AT2 cell differentiation in response to local signalling cues (Fig. 3J). Furthermore, the late-stalk cells are connected to AT2, AT1 and late airway progenitors in trajectory inference analysis (Fig. 2D; 3C,D), supporting our hypothesis that at all stages of lung development, cells exit the tip and enter a stalk-state prior to differentiation. This model for spatial patterning of human alveolar development is different from the current prevailing mouse model in which AT2 and AT1 cells are thought to be specified early in development (Zepp et al. 2021; Frank et al. 2019).

Integration of our fetal atlas with adult data revealed high correlation between expected groups: fetal airway progenitors with adult secretory club cells; fetal and adult ciliated and deuterosomal cells; proximal secretory fetal cells with adult goblet cells (Fig. S3A). The AT2 and AT1 cells we detect in the fetal lungs cluster closely with the adult (Fig. S3A; Pearson correlation coefficients: fetal-adult AT2 0.66; AT1 0.80). However, the fetal cells are immature and differ in gene expression to their adult counterparts (for example, for AT2/1, Fig. S3G).

### Lung endothelial cells exhibit early specialisation into arterial and venous identities

At 5-6 pcw, the endothelial cells (ECs) comprised capillary (early Cap: *THY1^+^, CD24^+^*), GRIA2^+^ arterial (*GRIA2^+^, GJA5^+^*) and lymphatic ECs (*PROX1^+^, STAB1^+^, UCP2^LO^*) (Fig. S8A-C); showing that capillaries and lymphatic vessels are distinct from the earliest stages of lung development and that arterial specification begins prior to venous. With increasing age, the capillary EC lineage moves from early (*THY1^+^, CD24^+^, EGLN1^+^*) to mid (*CA4^+^, KIT^+^, EGLN1^+^*) to late (*CA4^+^, KIT^+^*) sub-types (Fig. S8A-C), similar to the age transitions in other compartments. Trajectory analysis predicts that both mid- and late-Cap cells generate arterial and venous ECs (Fig. S8G,H). Aerocytes (*CA4^LO^, S100A3^+^*), capillary ECs specialised for gas exchange and leukocyte trafficking (Gillich et al. 2020; Vila Ellis et al. 2020), were observed at 20-22 pcw (Fig. S8A-C) arranged around the developing air sacs (Fig. S8D). Microvasculature specification therefore occurs relatively late in human fetal life coincident with the development of AT1 cells.

Broad markers of arterial and venous specification were clear in sections at 20 pcw (Fig. S8E). Three distinct arterial ECs were detected. GRIA2^+^ and arterial ECs (*DKK2^+^, SSUH2^+^*) form a continuous differentiation trajectory in pseudotime (Fig. S8G,H) with GRIA2^+^ ECs likely to be a more immature form. The OMD^+^ ECs (*GJA5^+^, DKK2^+^, PTGIS^+^, OMD^+^*) cluster with arterial ECs, are more proximal (Fig. S2H) and line the larger arterial vessels (Fig. S11B). By contrast, venous ECs (*PVLAP^+^, ACKR3^+^, HDAC9^+^*) do not have clear subclusters (Fig. S8A-C). Systemic and pulmonary circulation ECs have been found in adult lungs (Schupp et al. 2021); we cannot detect these in fetal lungs.

Two major lymphatic ECs were detected, lymphatic ECs (*PROX1^+^, STAB1^+^, UCP2^LO^*) and SCG3^+^ lymphatic ECs (*PROX1^+^, SCG3^+^*), with an intermediate population connecting them (Fig. S8A-C,F). SCG3^+^ lymphatic ECs resemble a lymphatic valve population (Takeda et al. 2019).

### Haematopoietic cell types in the developing lung

At the early stages (5-6 pcw) when arterial, capillary and lymphatic ECs were present, embryonic erythrocyte, HMOX1^+^ erythroblast and a small number of macrophages and ILC progenitors were detected, representing the early progenitors of haematopoiesis. After 11 pcw relative numbers of lymphoid and myeloid cells increased, dominated by macrophages, ILCs, dendritic, NK, T and B cells (Fig. 1C, S2A,B, S9A-C,F,G,K). Immature T cells are largely absent from the atlas, consistent with the restriction of T cell development to the thymus. In contrast, a range of early B cell precursors and the ILC precursor were detected. We enriched TCR and BCR fragments from our scRNA-seq libraries which supported cell-type identities and subdivision (Fig. S9D-E,H-J). To look for lung-specific features of the leukocyte cells, we compared our atlas with a pan-fetal human atlas (Cao et al. 2020). Unlike epithelial, endothelial and fibroblasts, leukocytes in our atlas are transcriptionally highly similar to those of other organs with minimal evidence of lung-specificity (Fig. S9L).

### Developmental trajectories of mesenchymal cells

The broad fibroblast cluster comprises fibroblasts, myofibroblasts, airway and vascular smooth muscle (ASM and vSMC), pericytes, mesothelium and chondrocytes (Fig. 4A,B). Airway fibroblasts and chondrocytes were proximally-enriched; mesothelium distally-enriched (Fig. S2H; 4D). There is a distinct separation of cell clusters by age (Fig. 4C). Airway SM cells were observed from 9 pcw, consistent with previous immunostaining (Nikolić et al. 2017), and showed increasing maturity over time (Fig. 4A-C). Two distinct populations of vSMC were observed throughout the time course: vSMC1 (*NTRK3^+^, NTN4^+^, PLN^-^*) and vSMC2 (*NTRK3^+^, NTN4^+^, PLN^+^*) (Fig. 4A,B) and were intermingled around the same vessels on tissue sections (Fig. S10A,C). vSMC1 was enriched in genes relating to ECM organisation and cell adhesion, whereas vSMC2 was enriched for transcripts encoding contractility proteins and signalling molecules (Fig. S10B). Intermingling of vSMC subtypes with different levels of contractility proteins is seen in adult lungs (Frid et al. 1997); our developmental observation suggests that these represent normal functional/ontological differences, rather than pathology. Pericytes (*FAM162B^+^*) were visualised adjacent to the microvascular endothelium (Fig. S10D).

**Figure 4.**
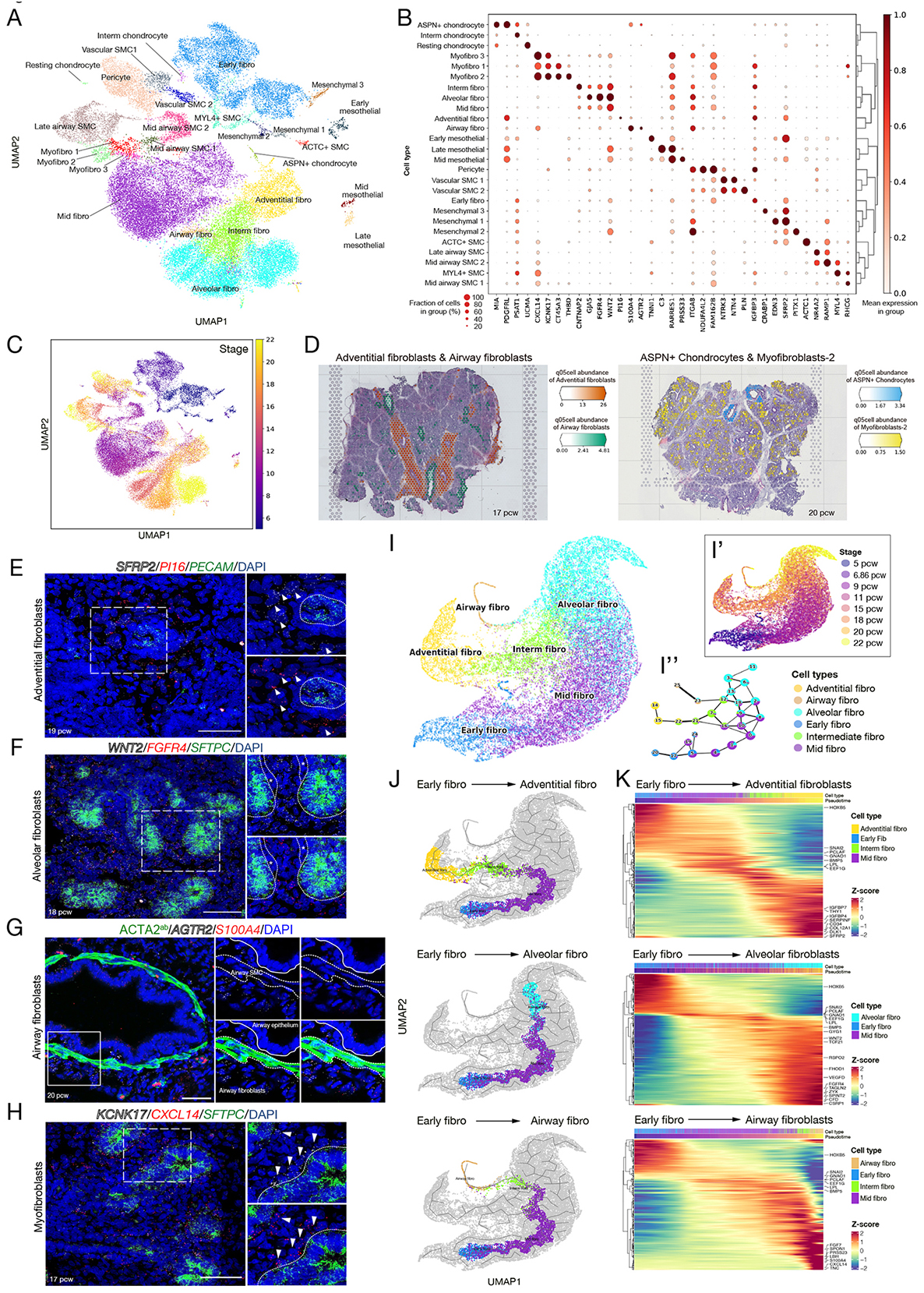
Diverse mesenchymal cell types localise to distinct niches in the developing human lung. (A) UMAP visualisation of mesenchymal cells. (B) Dot plot describing mesenchymal differential marker gene expression. (C) UMAP visualisation of mesenchymal cells coloured by stage. (D) Visium spatial feature plots visualising adventitial fibroblasts, airway fibroblasts, ASPN+ chondrocytes, and myofibroblasts-2 on 17 and 20 pcw lung sections. Scores are conservative estimates of cell-type abundance per voxel. (E-H) *In situ* HCR assay (E-H) and immunostaining (G). (E) Adventitial fibroblasts (*SFRP2,*white/*PI16,*red; arrowheads), ECs (*PECAM1*, green). (F) Alveolar fibroblasts (*WNT2* white; *FGFR4* red), tip cells (*SFTPC* green). Asterisks (* myofibroblasts). (G) Airway fibroblasts (*S100A4* red; *AGTR2* white), smooth muscle (ACTA2 green, dashed line). (H) Myofibroblasts (*KCNK17* white, *CXCL14* red; arrowheads), tip cells (*SFTPC*, green). DAPI, nuclei. Scale bars, 50 μm. (I) UMAP visualisation of cell types (I), stage (I’) and PAGA analysis (I’’) of fibroblast differentiation trajectories, . (J, K) UMAPs with Monocle3 trajectories (J) and selected trajectory gene expression heatmaps (K) for mid tip to adventitial fibroblasts (top), alveolar fibroblasts (middle), or airway fibroblasts (bottom). See also Figure S10.

The most common cells isolated from 5-15 pcw lungs were fibroblasts (Fig. 1C). At 5-6 pcw, early-fibroblasts (*SFRP2^+^,WNT2^+^*) predominated, although multiple populations were detected (Fig. 4A,B). In 9,11 pcw lungs, early-fibroblasts had matured into mid-fibroblasts (*WNT2^+^, FGFR4^LO^*) which have recently been shown to promote epithelial tip cell fate (Hein et al. 2022). In the most mature lungs sequenced, there were three distinct fibroblast populations: adventitial (*SFRP2^+^, PI16*^+^), airway (*AGTR2*^+^, *S100A4^+^*) and alveolar (*WNT2^+^, FGFR4^+^*) with distinct spatial locations (Fig. 4A,B,D-H). In addition, an intermediate fibroblast population connected the more mature fibroblasts on the UMAP (Fig. 4A,B), possibly representing a transitional state. Pseudotime analysis predicted a differentiation hierarchy from the early and mid fibroblasts to adventitial fibroblasts; with alveolar and airway fibroblasts forming separate branches (Fig. 4I-K). Alternatively, the intermediate fibroblast population may indicate plasticity in the fibroblast lineage in normal development as previously suggested (Kumar et al. 2014).

The three major fibroblast types in 15-22 pcw lungs expressed high levels of genes associated with ECM organisation, but had distinct gene expression patterns and spatial localisation. Adventitial fibroblasts (*SFRP2^+^, PI16^+^*) surrounded the larger blood vessels (Fig 4D). They formed diffusely arranged layers of cells surrounding the tightly packed concentric rings of ECs, pericytes and smooth muscle (Fig 4E, S10C). Adventitial fibroblasts were enriched in gene expression associated with ECM organisation and signalling, including BMP, TGFβ, WNT (Fig. 4J,K; S10E,F) consistent with described roles providing structural support to the perivascular region (Dahlgren and Molofsky 2019). Alveolar fibroblasts (*WNT2^+^, FGFR4^+^*) were observed throughout the lung, particularly surrounding the tip cells and close to the microvasculature (Fig 4F). They were enriched in genes associated with actin organisation, focal adhesions and morphogenesis, as well as signalling molecules (Fig. 4J,K; S10E,F). Adventitial and alveolar fibroblasts shared key markers such as collagens, but also expressed unique genes (adventitial: *SERPINF1, SFRP2, PI16*; alveolar: *FGFR4, VEGFD;* Fig. 4K). By contrast, the airway fibroblasts (*AGTR2*^+^, *S100A4^+^*, note that *S100A4* is expressed in various immune and airway epithelial cells) were adjacent to the airway smooth muscle and highly enriched in signalling molecules associated with morphogenesis (Fig. 4D,G,J,K; S10E,F). We did not detect lipofibroblasts (Travaglini et al. 2020), meaning that they are either exceptionally rare, or form later than 22 pcw, or do not form distinct clusters in all lung data sets (Madissoon et al. 2021). Endothelial and fibroblast populations align well between fetal and adult data (Fig. S3B,C), but with some unique developmental states, such as fetal early/mid-fibroblasts and myofibroblasts.

Myofibroblasts formed three distinct groups in our single cell data. Myofibroblast 1 (*CXCL14^+^, KCNK17^+^, CT45A3^+^, THBD^LO^*) appeared at 9 pcw and persisted to 20 pcw. Myofibroblast 2 (*CXCL14^+^, KCNK17^+^, CT45A3^+^, THBD^HI^*) and myofibroblast 3 (*CXCL14^+^, KCNK17^+^, CT45A3^-^, THBD^-^)* were predominantly identified at 22 pcw (Fig. 4A,B). Throughout development, myofibroblasts (*CXCL14^+^, KCNK17^+^*) were visualised surrounding the developing stalk region of the epithelium, suggesting a close signalling relationship (Fig. 4D,H, S10G,H). Although not detected in significant numbers in the scRNA-seq data until 22 pcw, we see myofibroblast 2 (PDGFRA^+^, THBD^HI^, *NOTUM^+^*) around the stalk epithelium from 15 pcw (Fig. 4D; S10H,J,K), the same position as myofibroblast 1. The appearance of myofibroblast 2 is coincident with the acquisition of AT2 markers by the late tip cells, and it may be a more mature state of myofibroblast 1. Myofibroblast 2 was enriched in gene expression associated with cell contractility and focal adhesions, as well as WNT signalling (Fig. S10J,K). Co-expression of the Wnt-responsive genes *LEF1*, *NOTUM* and *NKD1* suggests that myofibroblast 2 is responding to local Wnt expression (*WNT2* is high in alveolar fibroblasts) and producing the secreted Wnt inhibitor NOTUM; potentially to regulate local cell patterning. We tested this hypothesis using co-culture experiments where myofibroblast 2 was shown to both respond to WNT and to modulate the Wnt-response of co-cultured epithelial cells (Lim et al. 2021). *In vivo,* it is likely that myofibroblast 2 modulates the WNT2 signal from the alveolar fibroblasts mediating spatial patterning of epithelial AT2 and AT1 fate in human lung development. By contrast, myofibroblast 3 has higher expression of genes associated with ECM organisation and a variety of signalling molecules, including *C7*, *RSPO2* and *BMPER* (Fig. S10J). Myofibroblast 3 was always localised to the developing air sacs (Fig. S10I), rather than the stalk epithelium, and are likely to be precursors of the alveolar myofibroblasts (Li et al. 2020; Zepp et al. 2021).

### Signalling niches in lung development

We used CellPhoneDB (Efremova et al. 2020) to analyse cell-cell communication with the aim of predicting signalling interactions controlling cell fate allocation. We focused on 15-22 pcw cells and, based on the spatial localisation of the major fibroblast populations (Fig. 4E-G), analysed signalling within three niches, defined as - Airway niche: airway fibroblasts; late airway SMCs; airway epithelial cells. Alveolar niche: alveolar fibroblasts, aerocytes, late Cap cells, late tip cells, AT1, AT2. Adventitial niche: adventitial fibroblasts, arterial endothelium, OMD^+^ endothelium, vascular smooth muscle cells. CellPhoneDB predicts numerous signalling interactions (Supplemental Table 2) which we curated by plotting the expression of ligand-receptor pairs representing major signalling pathways (Fig. 5A,B; S11A). We observed expected interactions, including high levels of Notch ligands and receptors and CXCL12-CXCR4 signalling in the advential niche (Fig. S11A,B) (Herbert and Stainier 2011). Similarly, expected signalling predicted in the alveolar niche included aerocytes to late cap cells (ALPN-ALPNR) and alveolar epithelial cells to microvascular ECs (VEGFA-FLT1/FLT4/KDR) (Fig. 5B; S11B) (Gillich et al. 2020; Vila Ellis et al. 2020).

**Figure 5.**
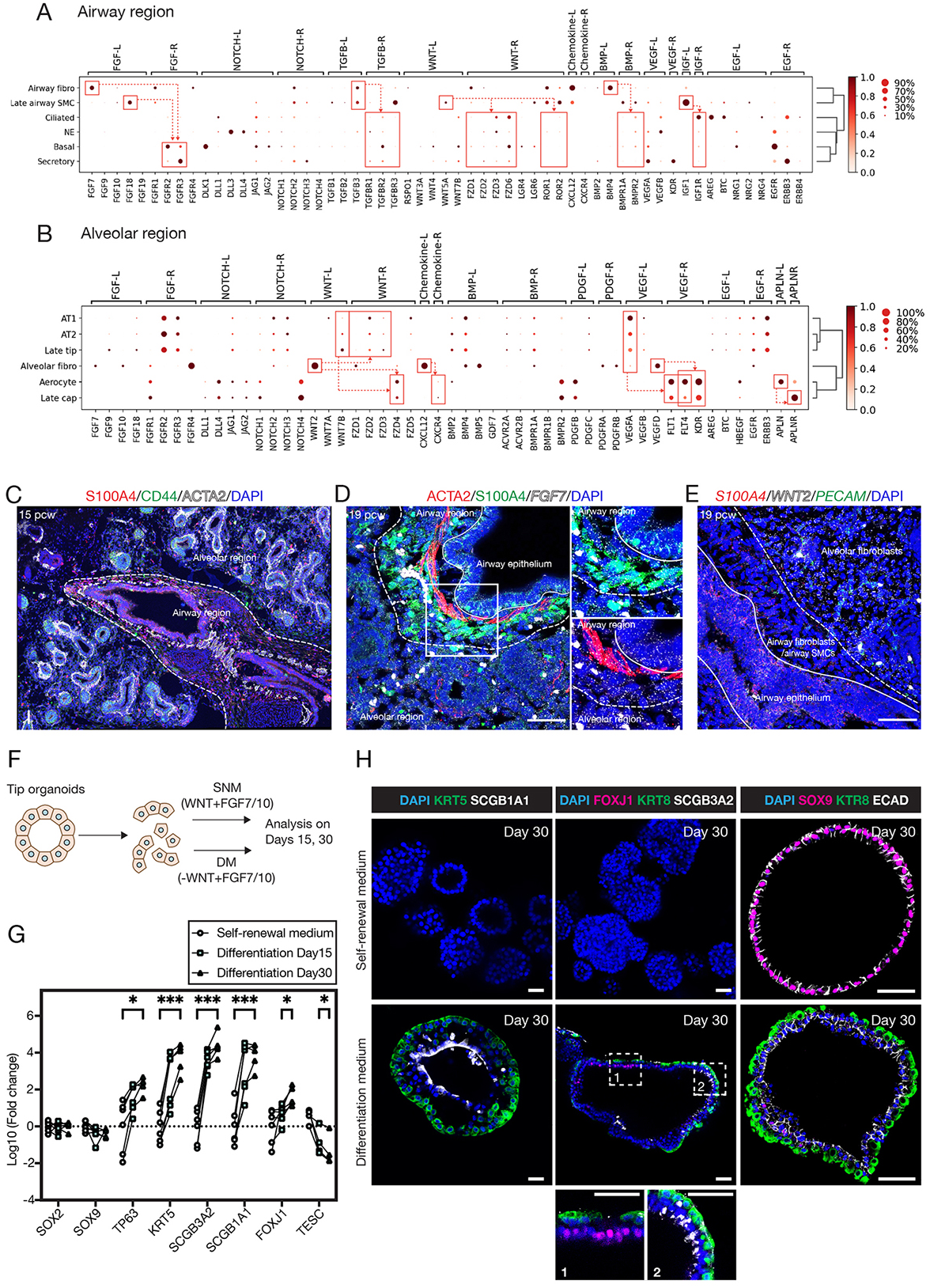
Signalling ligand-receptor interactions in alveolar and airway niches. (A, B) Curated ligand–receptor interaction predictions from CellPhoneDB in alveolar (A) and airway (B) niches. Dot plots visualise gene expression by cell type; dashed arrows indicate the predicted direction of signalling. (C-E) Immunofluorescence/HCR. S100A4/*S100A4*, airway fibroblasts; ACTA2, ASM; CD44, tip epithelium; PECAM1, ECs. Airway fibroblasts/ASM form a boundary (dashed lines) between alveolar and airway regions. Lines are between airway fibroblasts/SMCs and airway epithelium. DAPI, nuclei. Scale bars, 50 μm. (F) Organoids were cultured in FGF7/10-containing medium, in the presence (self-renewal medium; SNM) or absence (differentiation medium; DM) of CHIR99021, for 30 days. (G) qRT-PCR quantifications normalised to the organoids cultured in SNM. Significance evaluated by 2-way ANOVA with Tukey multiple comparison post-test; \**P*<0.05, \*\**P*<0.01, \*\*\**P*<0.001; *n*=6 organoid lines. (H) Whole mount immunofluorescence of lung organoids cultured in self-renewal medium (upper) and differentiation medium (lower). DAPI, nuclei. Scale bar, 25 μm. See also Figure S11.

Airway fibroblasts were predicted to signal via TGFβ3 and BMP4 to the airway epithelium, consistent with roles for these signals in human basal cell specification and differentiation (Miller et al. 2020; Mou et al. 2016). Airway fibroblasts and ASM were also predicted to signal to the epithelium via FGF7/18 to FGFR2/3 and non-canonical WNT5A to FZD/ROR (Fig. 5A). By contrast, although FGF and WNT signalling interactions were predicted in the alveolar niche, interactions were based on lower levels of *FGF*, but higher levels of canonical *WNT2* and its receptor (Fig. 5B). The predicted FGF and WNT signalling interactions in the alveolar niche/late tip cells are consistent with the requirement of these factors for long-term self-renewal of human distal tip organoids (Nikolić et al. 2017; Lim et al. 2021). Tissue staining showed that although *FGF7* is expressed fairly ubiquitously, the airway fibroblasts and ASM form a distinct barrier between the airway epithelium and the *WNT2* expression (Fig. 5C-E). Based on these data, we predicted that removing canonical WNT, but retaining FGF signalling would promote airway differentiation in the human distal tip organoids (Fig. 5F). Indeed, we observed robust basal, secretory and ciliated cell differentiation in response to FGF-containing medium (Fig. 5G,H).

### scATAC-seq analysis identifies putative cell fate regulators

Single cell ATAC-seq provides an independent method of assessing cell type based on open chromatin regions and allows cell type-specific TFs to be predicted. After tissue dissociation, the single cell suspensions were split and half of the cells processed for nuclear isolation and scATAC-seq (Fig. 1A). Following quality control and doublet removal, 67 scATAC-seq clusters comprising ∼100K cells were obtained and a label transfer process was used to annotate scATAC-seq clusters based on our scRNA-seq data (Fig. 6A). Not every cell state detected by scRNA-seq was distinguishable by scATAC-seq, consistent with previous work (Domcke et al. 2020; Cao et al. 2020). For example, separate early-tip, stalk and airway progenitor clusters were discerned by scRNA-seq (Fig. 2A), but a combined cluster with strong similarity to all three cell types was detected by scATAC-seq (Fig. 6A). Similarly, the resolution of scATAC-seq allowed us to identify a combined AT1/AT2 cluster and single arterial endothelial, vascular smooth muscle, myofibroblast and basal cell clusters (Fig. 6A). Nevertheless, there was broad agreement between the scRNA-seq and ATAC-seq data in terms of capturing cell types, including many of the novel/lesser-known cell types we identified by scRNA-seq (mid and late tip, mid and late airway progenitors, GHRL^+^ NE, MUC16^+^ ciliated, dueterosomal, airway fibroblasts, aerocytes, SCG3^+^ lymphatic endothelial cells).

**Figure 6.**
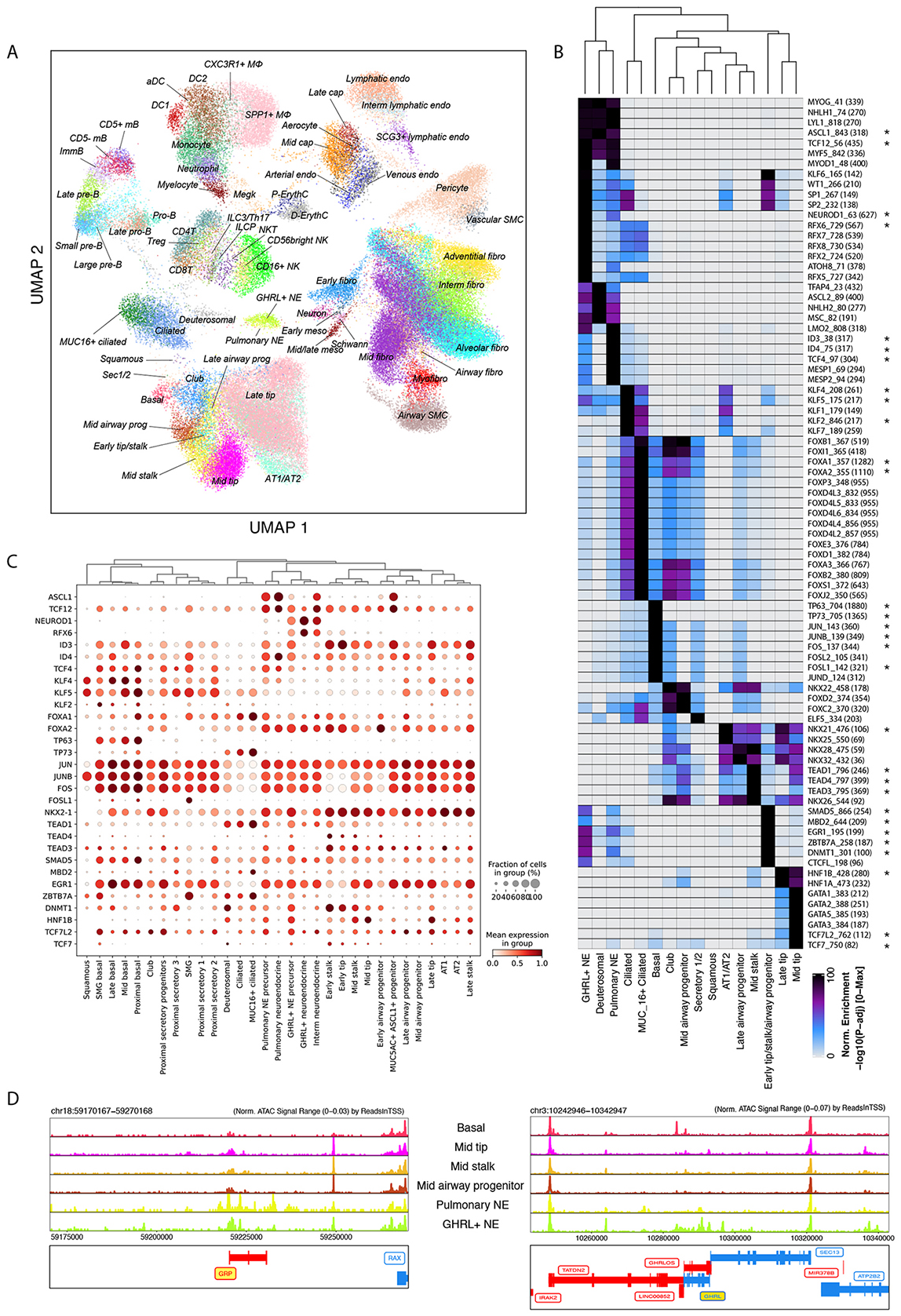
DNA accessibility and motif enrichment revealed by scATAC-seq. (A) Single-cell DNA accessibility profiles mapped onto 2D UMAP plane. Coloured for cell states. (B) A union set of top 10 enriched motifs in the marker peaks among epithelial cell types/states. Statistical significance is visualised as a heatmap according to the colour bar below. Transcription factors concordantly expressed based on scRNA-seq data are marked with asterisks. (C) Expression dotplot of the concordant transcription factors from (B) in epithelial cell types. (D-E) Read coverage tracks of in silico aggregated “pseudo-bulk” epithelial clusters over the GRP locus (D) and GHRL locus (E). See also Figure S12.

We analysed TF binding motifs in the unique/enriched open chromatin regions in each cluster and plotted the top 5 TF motifs per cell type (Fig. S12). As expected, TFs belonging to the same family are frequently enriched in the same cell type due to similarities in their binding motifs. This analysis revealed some expected TF signatures, for example TCF21 in the alveolar, adventitial and airway fibroblasts (Quaggin et al. 1999), GRHL and FOXA1/2 in epithelium (Gao et al. 2013; Wan et al. 2005), and SOX17 in arterial endothelium (Corada et al. 2013). Examining epithelial cells and focussing on TFs expressed in the corresponding cell type in the scRNA-seq data (Fig. 6B,C, marked by asterisk in B), TEAD motifs were enriched in mid-stalk cells consistent with a key role for Yap (van Soldt et al. 2019), NKX2.1 in AT1/AT2 cells (Kimura, Ostrin, and Chen 2019), KLF factors in secretory and AT1/AT2 (Liberti et al. 2022) and TP63 in basal cells (Rock et al. 2009). Unexpected TF signatures included HNF1B in late tip cells, and ZBTB7A in early tip/stalk/airway progenitors. We focused on the pulmonary and GHRL^+^ NE cells which cluster closely in both data sets (Fig. 2A, 6A). ASCL1 is required for mouse NE cell differentiation (Ito et al. 2000; Borges et al. 1997) and this motif is strongly associated with both pulmonary and GHRL^+^ NE cells (Fig. 6B). However, both cell types also have specific TF motifs including NEUROD1 and RFX6 in the GHRL^+^ NEs, and TCF4 and ID in the pulmonary NEs (Fig. 6B). Consistent with this, there are distinct, unique regions of open chromatin, especially in the neighbourhood of cell-type specific genes such as GRP and GHRL (Fig. 6D).

We have produced a high-resolution scATAC-seq data set for the developing human lungs which is highly consistent with our scRNA-seq data. Mining this data provides hypotheses for lineage-determining TFs in lung development. As further scATAC datasets become available for control and diseased adult lungs, our data will provide a resource for comparing normal and aberrant TFs and chromatin regulation.

### Transcriptional control of neuroendocrine cell subtype formation

Pulmonary NE and GHRL^+^ NE cells share the expression of many TFs and open chromatin regions, but are transcriptionally distinct. In our scRNA-seq data, they were both observed along a maturation trajectory (from what are labelled as precursors), shared classical NE markers (*CHGA, SYP*), but differed in TF and hormone expression (Fig. 7A,B). A third NE population (intermediate NE) clustered between pulmonary and GHRL^+^ NE cells with intermediate gene expression (Fig. 7A,B), although it did contain a small number of cells expressing the unique marker *NEUROG3.* Pseudotime trajectory analysis suggested that pulmonary NE and GHRL*^+^* NE cells were derived from airway progenitors/stalk cells and that intermediate NEs are an additional transition population (Fig. S13A,B). Transition states between pulmonary NE and GHRL^+^ NE were observed in sections (Fig. S13C). We therefore postulated that Pulmonary NE precursors could acquire *NEUROG3* and convert to GHRL^+^ NE fate (Fig. 7C), or vice-versa - GHRL^+^ precursors converting to pulmonary NE fate. In sections, *ASCL1* was co-expressed with *GRP*, but rarely with *GHRL*. We also observed *ASCL1* single-positive cells, likely representing pulmonary NE precursors (Fig. 7D). *NEUROD1* was co-expressed with *GHRL*, but also observed with *GRP* (Fig. 7E). Whereas *NEUROG3* was co-expressed with *ASCL1* and/or *NEUROD1*, supporting a role in a transition population (Fig. S13D).

**Figure 7.**
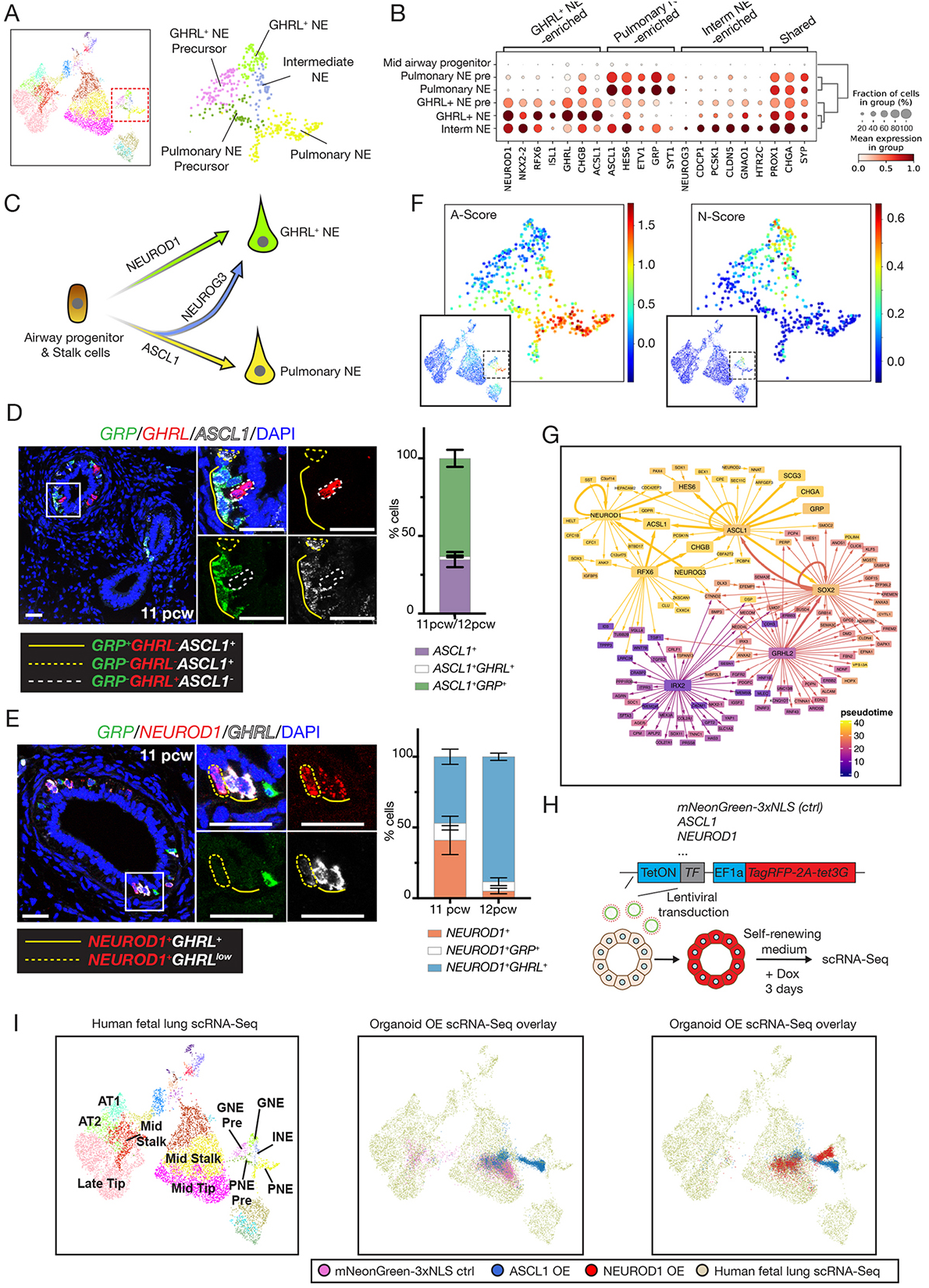
*ASCL1* and *NEUROD1* regulate the formation of two subtypes of neuroendocrine cells. (A) Zoom-in UMAP plot of NE lineages. (B) Dot plot showing selected gene expression in NE lineages. (C) Schematic model of NE lineage formation. (D) Left: HCR, *GRP* (green), *GHRL* (red), *ASCL1* (white). Right: Mean ± SEM of *ASCL1*^+^ cell types, N = 3 human fetal lungs, n = 243 *ASCL1*^+^ cells. (E) Left: HCR, *GRP* (green), *NEUROD1* (red), *GHRL* (white). Right: Mean ± SEM of *NEUROD1*^+^ cell types: N = 2, 11 pcw human fetal lungs, n = 129; N = 3, 12 pcw human fetal lungs, n = 132. Scale bars 25 μm. (F) Gene signature scoring of A-type and N-type SCLC features in the epithelial UMAP. (G) Scenic analysis of predicted TF network governing mid tip progenitor cells to pulmonary NE. Trajectory and colour coding match Fig. S13A,B. (H) Organoids from 8 pcw human fetal lungs were transduced with Doxycline (Dox) inducible TF, or mNeonGreen-NLS, lentivirus. Transduced organoids were isolated by flow cytometry based on TagRFP expression, seeded in Matrigel for 10-13 days prior to Dox treatment. Organoid cells were harvested 3 days post-Dox for scRNA-Seq. N=3 organoid lines. (I) Left: reference UMAP of primary human fetal lung epithelium. Mid and right: scRNA-Seq of organoids overexpressing *mNeonGreen-NLS*, *ASCL1* or *NEUROD1* projected onto the primary data. See also Figures S13 and S14.

Differential expression of *ASCL1* and *NEUROD1* defines A- and N-type human small cell lung cancer (SCLC), which likely derive from NE cells (Gay et al. 2021). Interestingly, these two TFs coincide with the scRNA-seq marker genes and scATAC-seq TF motif enrichment of our fetal NE cells (Fig. 6B,7B). We generated SCLC feature gene lists (Borromeo et al. 2016) and performed gene signature scoring, showing that the A-type signature resembles pulmonary NEs, whereas the N-type resembles GHRL^+^ NEs (Fig. 7F). These data suggest that either there are two different NE cells of origin for human SCLCs, or that SCLCs reuse developmental mechanisms as suggested by some mouse models (Ireland et al. 2020). We have been unable to detect GHRL^+^ NEs in the adult airways using HCR (5 biological replicates). However, a small number of GHRL^+^ cells are present within a tuft cell cluster in an integrated adult lung cell atlas containing 2.2 million cells (Sikkema et al. 2022), suggesting that GHRL^+^ NEs could be a rare cell state in the adult airways. Given their relevance to human disease states, we sought to use our single cell atlas to identify lineage-defining TFs controlling NE cell differentiation and test these predictions using our organoid system. We reasoned that overexpression of lineage-defining transcription factors in lung tip organoids (Nikolić et al. 2017; Sun et al. 2021) would promote cell type-specific differentiation.

Multiple TFs were differentially expressed between pulmonary NE and *GHRL^+^* NE cells (Fig. 7B). We used SCENIC analysis of gene regulatory networks (GRNs) (Aibar et al. 2017) along a predicted airway progenitor to GHRL^+^ NE trajectory (Fig. S13A,B) to identify putative lineage-defining TFs (Fig. 7G). ASCL1, NEUROD1 and NEUROG3 all emerged as potential key nodes and are required for endocrine cell differentiation in various organs (Borges et al. 1997; Ito et al. 2000; Mellitzer et al. 2010; Naya et al. 1997). We also selected the *GHRL^+^*NE-specific *RFX6* (Fig. S13E) and *NKX2.2* (Fig. 7B), the pan-NE *PROX1* (Fig. 7B) and, as controls, the basal cell-specific TFs *DeltaNTP63*, *TFAP2A*, *PAX9,* and *mNeonGreen-3xNLS*. Overexpression of *PROX1* or *NKX2-2* did not result in NE gene upregulation based on qRT-PCR (not shown) and these TFs were not followed up. The other factors resulted in increased expression of basal or NE markers compared to *mNeonGreen-3xNLS* controls and the experiments were repeated using scRNA-seq. Individual TFs were overexpressed from a doxycycline-inducible construct for 3 days and organoids were maintained in the self-renewing (tip cell-promoting) medium throughout to rigorously assay the lineage-determining competence of the TF (Fig. 7H; S14A), followed by scRNA-seq.

When mapped to epithelial cells of our fetal lung atlas, the majority of the *mNeonGreen-3xNLS* expressing organoid cells projected to mid-tip or -stalk cells as expected (Fig. 7I). Whereas, overexpression of *DeltaNTP63* resulted in basal cell-like lineages (S14B) consistent with a previous report (Warner et al. 2013), indicating that this simple assay can report TF function. Overexpression of *RFX6*, *TFAP2A* or *PAX9* did not result in the predicted lineage progression at a transcriptome level (S14B). However, *ASCL1*-overexpressing organoids progressed into pulmonary NE precursors (Fig. 7I) and *NEUROD1* overexpression promoted differentiation into *GHRL*^+^ NE precursors (Fig. 7I). *NEUROG3* overexpression also led to *GHRL*^+^ NE precursor formation (Fig. S14B), suggesting that the *GHRL^+^* NE lineage is the destination of the intermediate NE population (Fig. 7C).

The 5’ differences between the transgenes and endogenous TFs allowed us to distinguish these transcripts and infer gene regulation hierarchy. We observed autoregulation of *ASCL1*, *NEUROD1*, *NEUROG3* and *RFX6* (Fig. S14C). By contrast, *NKX2-2* and *PROX1* were upregulated by other TFs, indicating they are relatively low in the hierarchy (Fig. S14C). *NKX2-2* and *PROX1* expression in the organoid assay matched their expression in NE cells *in vivo* (Fig. 7B, S14C), showing that this assay recapitulated key features of the TF network. These experiments have allowed us to test gene GRN predictions from the single cell atlas, confirm the predicted lineage trajectory and provide a foundation for studying human SCLC. This is significant given that there is no evidence that GHRL^+^ NE cells are present in mice (Borromeo et al. 2016), making the use of mouse models difficult.

## Discussion

Using a combination of single cell and spatial approaches we have identified 144 cell types, or states, in the developing human lungs across the 5-22 pcw period. We take advantage of a known proximal-distal gradient in epithelial differentiation to identify progenitor and differentiating states in the developing airway, including a new neuroendocrine cell subtype related to SCLC. We suggest that human alveolar epithelial differentiation follows a tip-stalk-AT2/1 fate decision pattern that is different to the prevailing cellular models of mouse alveolar development. Moreover, analysis of the mesenchymal compartment identified three niche regions with distinct signalling interactions, allowing us to identify signalling conditions that are sufficient for airway differentiation of human embryonic lung organoids. We tested GRN predictions for NE cell differentiation in an organoid system, allowing us to identify lineage-defining TFs and provide directionality to the inferred differentiation trajectory. This study provides a paradigm for combining single cell datasets with spatial analysis of the tissue and functional analyses in a human organoid system to provide mechanistic insights into human development.

We show that lung maturation occurs in concert across cell compartments, for example epithelial AT1 cells and endothelial aerocytes differentiating in parallel at 20-22 pcw. Moreover, we have observed many aspects of cellular differentiation *in utero* showing that they are controlled by prenatal factors, rather than the transition to air breathing with its associated mechanical and hormonal changes. For example, two distinct types of vSMCs and lymphatic endothelial cells are established prior to major alterations in blood flow that occur postnatally.

The mesenchymal compartment contains multiple fibroblast and myofibroblast cell states and we focus on those present during the later stages of lung development. Airway, adventitial and alveolar fibroblasts are all localised in distinct niche regions and participate in different signalling interactions. Airway and adventitial fibroblasts both express unique combinations of signalling molecules and also form physical barriers between the neighbouring airway epithelium, or vascular endothelium, and the widespread alveolar fibroblasts (Fig. 4,5). Similarly, we characterise a population of myofibroblasts which contacts the developing epithelial stalk region and expresses high levels of the secreted Wnt-inhibitor, *NOTUM* (Fig. S10K); whereas alveolar fibroblasts express high levels of the canonical *WNT2* ligand (Fig. 4). In a separate study, using surface markers identified in this single cell atlas, we were able to specifically isolate alveolar fibroblasts and myofibroblast 2 cells and perform co-culture experiments with late tip organoids (Lim et al. 2021). Those experiments confirmed that a three-way signalling interaction between alveolar fibroblasts, myofibroblast 2 cells and late tip cells can control human AT2 spatial patterning.

We find that GHRL^+^ NE cells are transcriptionally similar to the NEUROD1^+^ N-subtype of SCLC (Fig. 7). Our functional analyses of NE cell differentiation in organoids will provide tools to test these hypotheses. Mouse studies show that fetal transcriptional and chromatin cell states are accessed during the normal process of tissue regeneration and may contribute to neoplasm in chronic inflammation (Larsen and Jensen 2021; Jadhav et al. 2017). Detailed ATAC-seq datasets are not yet available for human lung disease. Our high quality ATAC-seq atlas will provide a baseline for further analyses when adult chromatin accessibility lung atlases are published. In summary, our multi-component atlas is a community resource for future analyses of human development, regeneration and disease.

### Limitations of the study

We provide a carefully annotated, descriptive cell atlas resource. Many conclusions are derived from trajectory inference or TF binding site analyses and require future validation. The trajectory inference analyses predict lineage relationships and transitions between cell types. However, these transitions reflect similarities in gene expression, for which direct differentiation processes are only one possible explanation. Further, they do not imply directionality. RNA velocity (used in Fig. 2D) provides directionality to trajectory inference predictions using the ratio of spliced versus unspliced mRNA and assuming steady-state transcription and degradation rates. However, over developmental time these assumptions are unlikely to hold across all developmental transitions, which can lead to the inference of incorrect directionality (Bergen et al. 2021). Similarly, when applying RNA velocity algorithms to scRNA-seq data of a known differentiation trajectory, reversed velocities have been reported (Gorin et al. 2022). For these reasons, we performed most of our trajectory inference analysis using Monocle3 (Trapnell et al. 2014). Monocle3 requires user-defined starting and end points and calculates the most likely routes between these points (shown as grey lines on the plots), guided by known biological features of the data (age and spatial arrangement of cells). A further confounding factor for trajectory inference in development is that upon maturation, some cell types are likely to act as progenitors themselves (Rawlins, Okubo, et al. 2009; Rock et al. 2009; Yang et al. 2018). Adding to the difficulty of reconstructing complete lineages, our single-cell dissociation protocols are expected to under-sample certain cells, such as mature neurons and chondrocytes. Furthermore, validation assays for lineage analysis in human systems rely on *in vitro* experiments, including organoid and iPSC differentiation. It is important to acknowledge that these usually define differentiation competence and do not necessarily mean that a specific differentiation route occurs *in vivo*.

We have performed data integration and regression analysis to compare the identity of our fetal human lung cells with adult human lung cells. There are approximately three decades between the oldest fetal and youngest adult human lung samples sequenced, including a rapid period of postnatal growth and morphogenesis, puberty and an unknown number of infections/environmental insults. Despite this, many fetal-adult similarities can be seen. Nevertheless, it will be important to sequence additional lungs and, when possible, to fill the age gap. Similarly, mouse-human fetal lung cell comparisons have discerned similarities and differences. However, the differences in experimental protocols and annotation granularity between the mouse and human data might have also contributed to the differences we see. Moreover, mice were selected as lab animals partly due to their small size and rapid gestation. It will be informative in the future to make comparisons with a range of fetal lungs, including larger, long-developing species such as pig and sheep, to distinguish between differences due to species, size and gestation period.

## Supporting information

Supplementary Table 1 - Cell type markers

Supplementary Table 2 - CellphoneDB

Supplementary Table 3 - Sample metadata

Supplementary Table 4 - HCR probes

Supplementary Table 5 - ATAC-seq motifs

## Acknowledgements

We would like to acknowledge the Gurdon Institute Imaging Facility, and the Cellular Genetics IT and Phenotyping group, New Pipeline Group and DNA pipelines of Sanger Institute; Menna Clatworthy and Muzz Haniffa and RE, CS, ED, IG, MH, CD, WS for discussions on cell-type annotations; MP, AP, SL and CT for informatics support. KL is supported by the Basic Science Research Program through the National Research Foundation of Korea (NRF) funded by the Ministry of Education (2018R1A6A3A03012122). DS is supported by a Wellcome Trust PhD studentship (109146/Z/15/Z) and the Department of Pathology, University of Cambridge. PH holds a non-stipendiary research fellowship at St Edmund’s College, University of Cambridge. EM is supported by ESPOD fellowship of EMBL-EBI and Sanger Institute. JPP is supported by the MSCA Postdoctoral Fellowship. ELR is supported by the MRC (MR/P009581/1; MR/S035907/1). ZD is supported by a Wellcome Trust PhD studentship (222275/Z/20/Z). RAB is supported by the NIHR Cambridge Biomedical Research Centre (BRC-1215-20014) and was an NIHR senior investigator. Gurdon Institute Core support from the Wellcome Trust (203144/Z/16/Z) and Cancer Research UK (C6946/A24843). KBM, JCM and SAT acknowledge funding from the MRC (MR/S035907/1) and from Wellcome (WT211276/Z/18/Z and Sanger core grant WT206194). MZN acknowledges funding from a MRC Clinician Scientist Fellowship (MR/W00111X/1), the Rosetrees Trust (M899) and Action Medical Research (GN2911). This work was partly undertaken at UCLH/UCL who received a proportion of funding from the Department of Health’s NIHR Biomedical Research Centre’s funding scheme.

## Author Contributions

Conceptualization: PH, KL, DS, KBM and ELR. Methodology: PH, KL, DS. Software: PH. Formal Analysis: PH, JPP, KP, ZKT. Investigation: KL, DS, QJ, ZD, LB, LR, LM, MD, AW and MY. Resources: EM, XH, RAB, SMJ. Data Curation: PH, KL, DS, EM, ZKT, ED, CS, IG. Writing-original draft: PH, KL, DS, JPP, KBM and ELR. Writing - review and editing: PH, KL, DS, KBM, ELR, SAT, JBM. Supervision: MZN, RAB, SAT, JBM, KBM and ELR. Funding Acquisition: KL, EM, JPP, RAB, MZN, SAT, JBM, KBM and ELR.

## FIGURE LEGENDS

**Figure S1.**
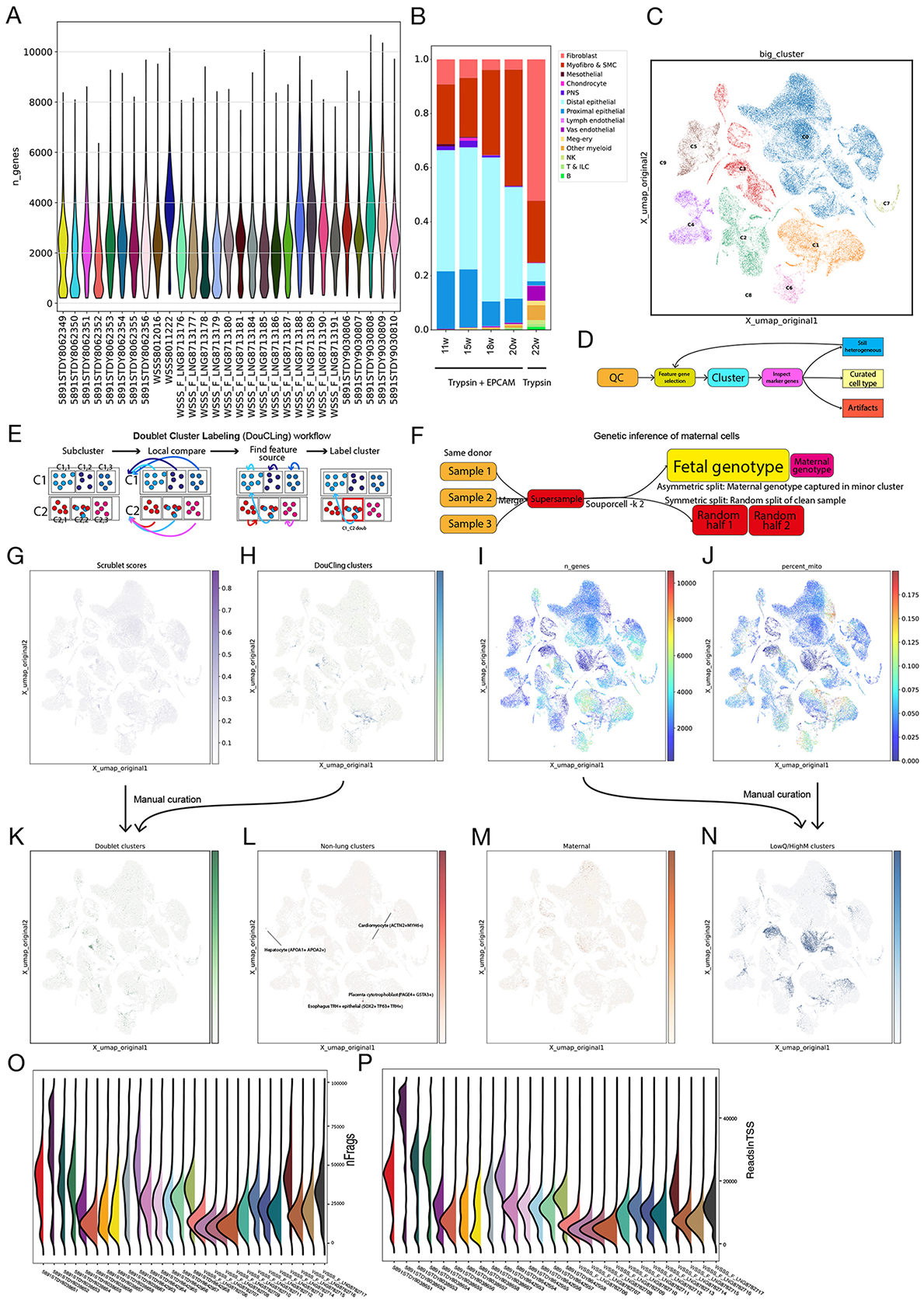
Quality control for scRNA-seq and scATAC-seq data. (A) Distributions of the number of genes detected per cell, grouped by 10X libraries. (B) Proportions of broad cell types in samples treated with Trypsin, and Trypsin plus EPCAM enrichment following colour codes in Figure 1C. (C) Initial clusters of data separating compartments, before subclustering. (D-F) Workflows of the recursive subclustering method (D), the Doublet Cluster Labeling (DouCLing) method to identify doublet-driven clusters (E), and inference of maternal cells using Souporcell (F). (G) Doublet scores calculated by Scrublet. (H) Inferred doublet clusters using DouCLing. (I) Number of genes detected projected on UMAP. (J) Percentage of mitochondrial reads. (K) Cells in curated doublet clusters. (L) Cells in clusters of cells coming from other organs. Marker genes in parentheses. (M) Inferred maternal cells. (N) Cells in curated low-quality cell clusters. (O,P) scATAC-seq quality metrics of fragment detection per cell (O) and reads mapped in transcription-start sites (P).

**Figure S2.**
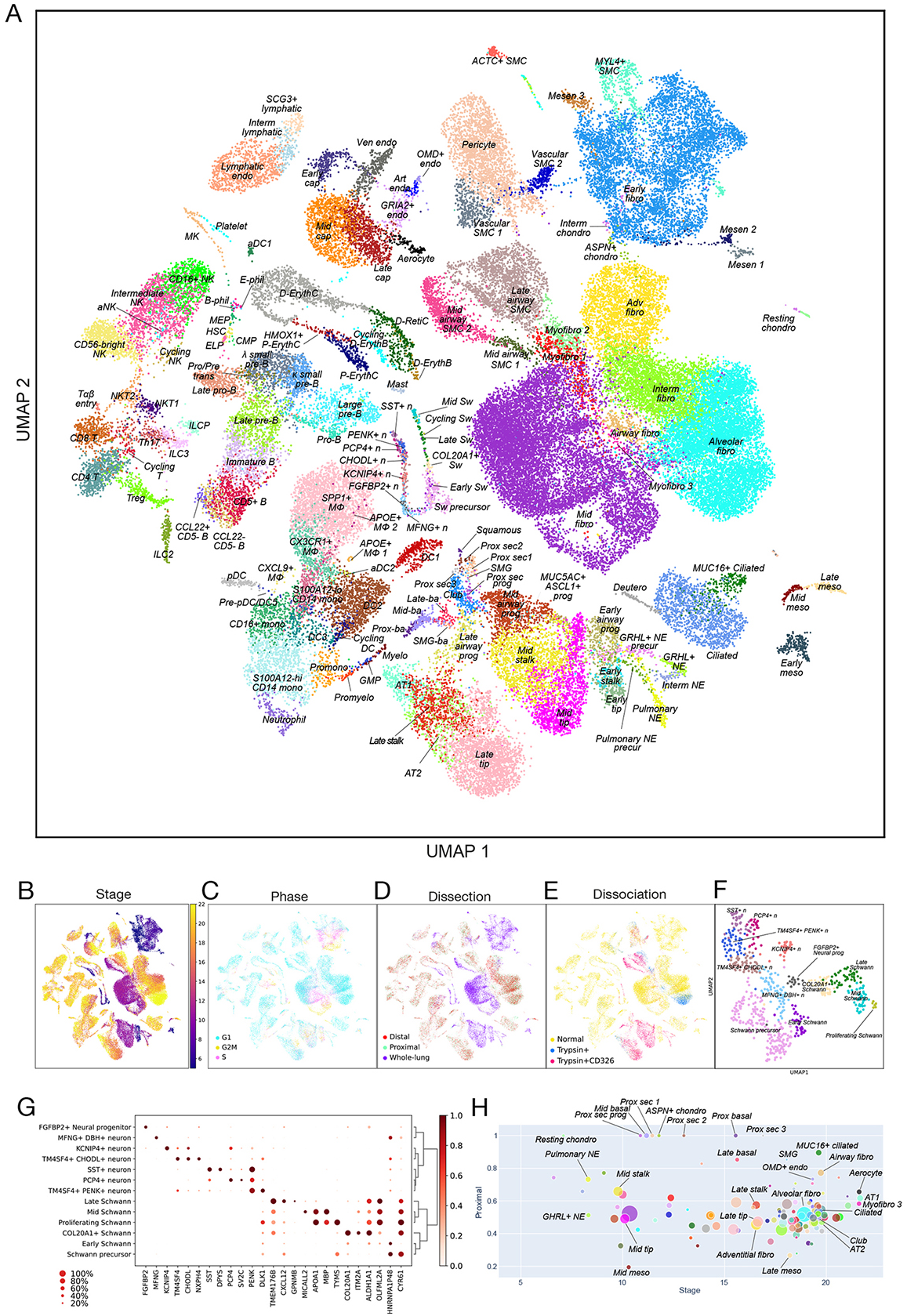
Overview of 144 cell types or cell states. (A-E) All of the curated 144 clusters of single cells projected on UMAP space of transcriptomes, colored by cell type/state (A), developmental stage (B), inferred cell-cycle phase (C), dissection region (D) and dissociation/enrichment strategy (E). (F) Cells from the initial PNS cluster (C7) projected on UMAP space of transcriptomes, colored by cell type/state. (G) Selected feature genes of cell types/states in the initial PNS cluster. (H) Spatiotemporal biases of cell types. Cell types are shown as dots with x representing the weighted average of developmental stages, y representing the score of proximal enrichment and the size corresponding to the cluster size.

**Figure S3.**
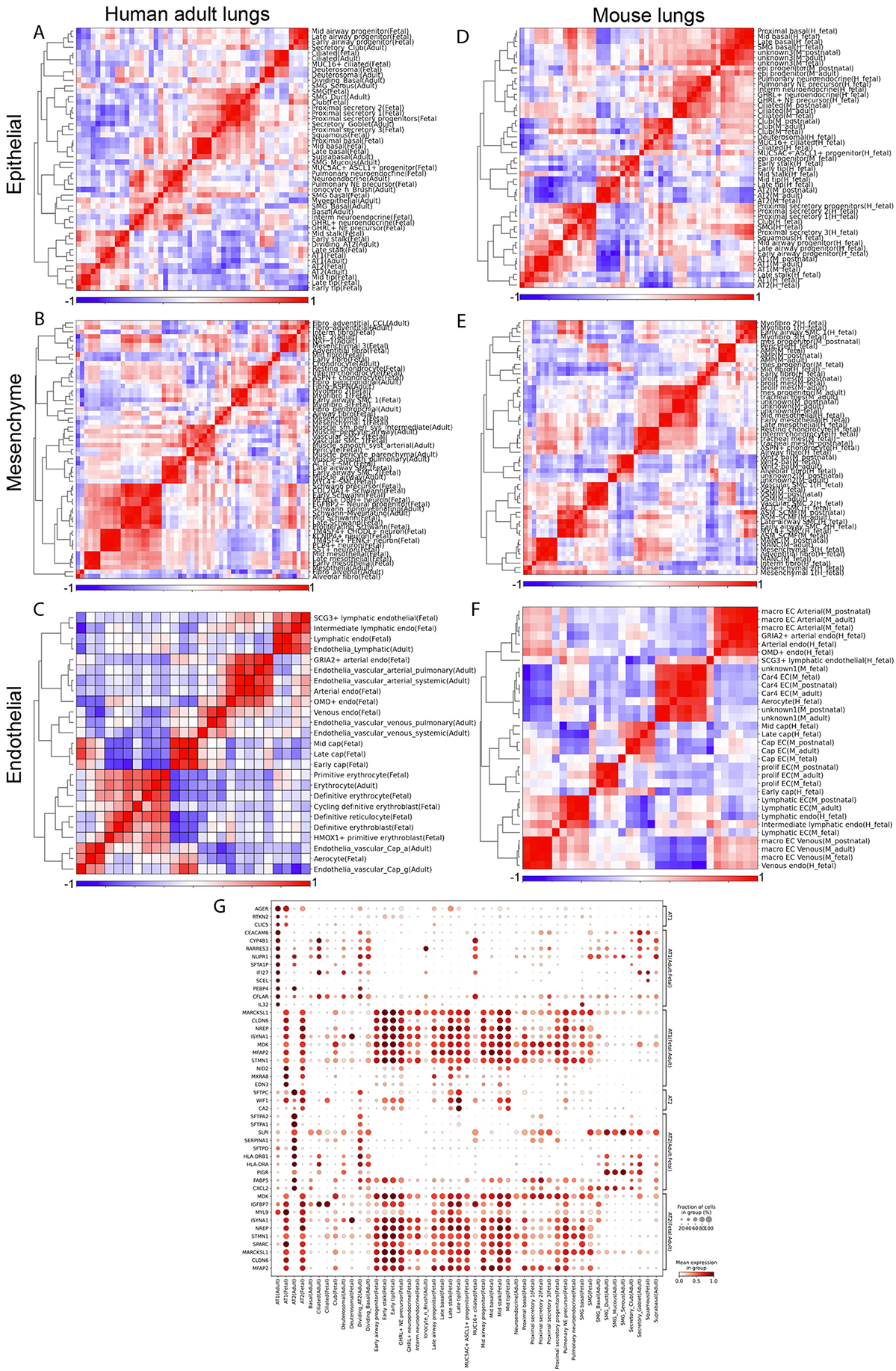
Comparing fetal lung scRNA-seq with adult human and mouse lung scRNA-seq. (A-F) Correlations of scVI latent variables between human fetal lung cell clusters and those of previously annotated adult cell clusters (Madissoon et al. 2021, A-C) and mouse lung cell clusters (Zepp et al. 2021, D-F), focusing on epithelial (A, D), fibroblast (B, E) and endothelial (C, F) compartments. (G) Expression dotplot of genes shared or unique to fetal/adult lung AT1/AT2 cell clusters..

**Figure S4.**
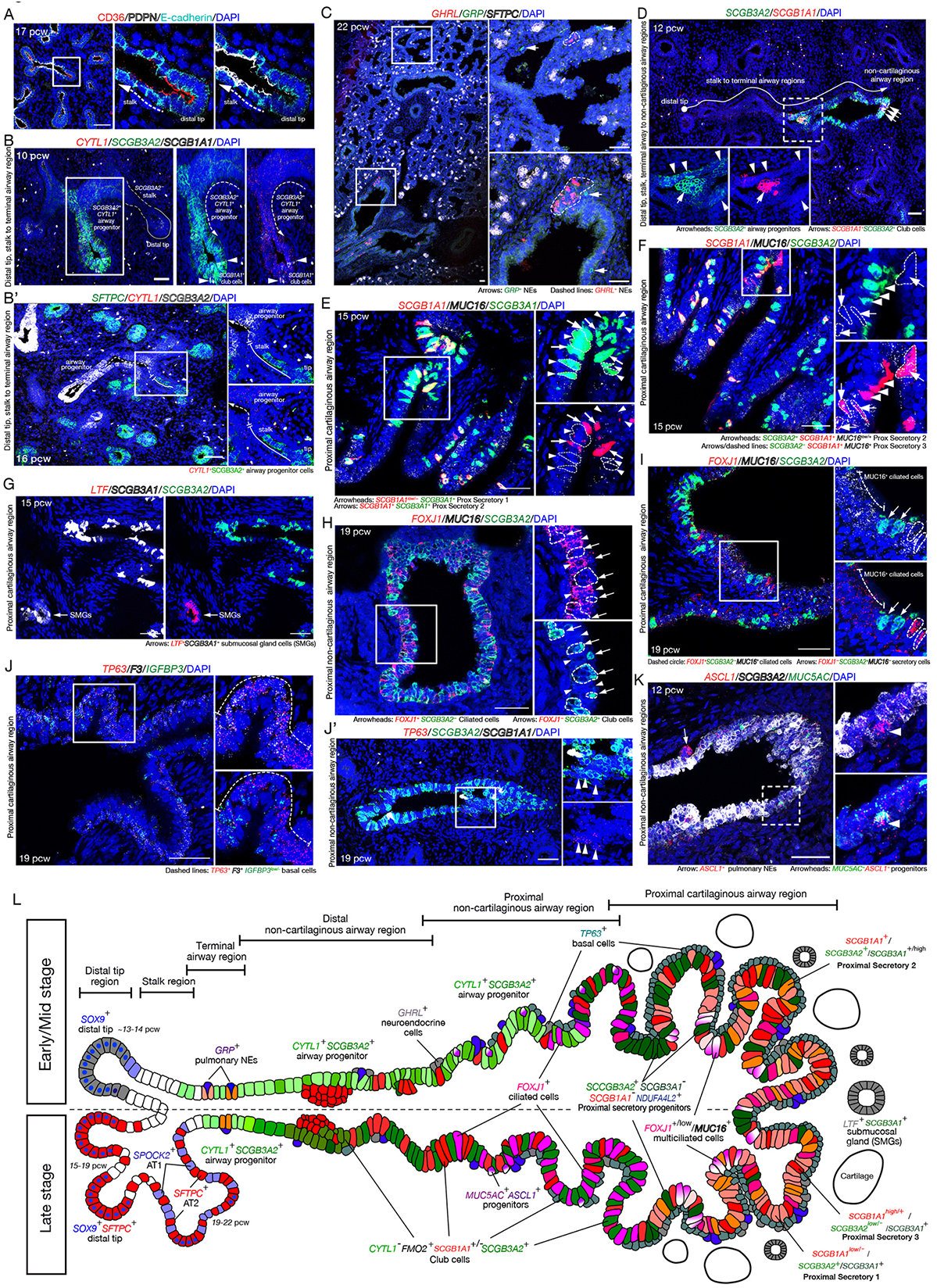
Spatial analysis of airway epithelial cells in the developing human lungs by *in situ* HCR. (A) Tip and stalk epithelial cells in distal regions of fetal lungs at 17 pcw, immunostained using antibodies against CD36 (tip epithelial cells, red), PDPN (stalk epithelial cells, white), and E-cadherin (epithelium, cyan). (B, B’) Airway progenitor cells in distal fetal lungs at 10 (B) and 16 (B’) pcw. The airway progenitor cells marked by *SOX9^-^/CYTL1^+^/SCGB3A2^+^* are located proximally to the *CYTL1^-^/SCGB3A2^-^* stalk. *SCGB1A1* indicates club cells (B, white). *SFTPC* is mainly expressed in the tip and partly located in stalk regions (B’, green). (C) GHRL^+^ neuroendocrine (dashed line, red) and GRP^+^ pulmonary neuroendocrine cells (arrow, green) in fetal lungs at 22 pcw. *SFTPC* indicates tip epithelial cells (white). (D) Airway progenitor (arrowhead) and club cells (arrow) in non-cartilaginous airway regions of fetal lungs at 12 pcw are marked by *SCGB3A2^+^/SCGB1A1^-^*and *SCGB3A2^+^/SCGB1A1^+^*, respectively. Tip, stalk, airway progenitor, and club cells are localised progressively more proximally from the distal tip regions to the proximal non-cartilaginous airway regions. *SCGB3A2* (green), *SCGB1A1* (red). (E) Proximal secretory 1 (arrowhead) and 2 (arrow) are distinguishable by the presence or absence of *SCGB1A1* expression, each marked by *SCGB3A1*^+^/*SCGB1A1^low^*^/-^/MUC16^low/-^ and *SCGB3A1*^+^/*SCGB1A1*^+^/MUC16^low/+^, respectively, in the proximal cartilaginous airway in 15 pcw fetal lungs. *MUC16*^+^ only cells are MUC16^+^ ciliated cells. *SCGB3A2* (green), *SCGB1A1* (red), *MUC16* (white). (F) Proximal secretory 2 (arrowhead) and 3 (arrow) are distinguishable by the presence or absence of *SCGB3A2* and *MUC16* expression, marked by *SCGB3A2*^+^/*SCGB1A1*^+^/MUC16^low/+^ and *SCGB3A2*^low/-^/*SCGB1A1^+^*/MUC16^+^, respectively, in the proximal cartilaginous airway of fetal lungs at 15 pcw. *SCGB3A2* (green), *SCGB1A1* (red), *MUC16* (white). (G) Submucosal gland cells (arrow) located in SMGs are marked by strong *LTF* expression with *SCGB3A1*^+^/*SCGB3A2*^-^ in the proximal cartilaginous airway regions of fetal lungs at 15 pcw. *SCGB3A2* (green), *LTF* (red), *SCGB3A1* (white). (H) Ciliated cells and secretory cells are distinguishable by expression of *FOXJ1* (red) or *SCGB3A2* (green) in the non-cartilaginous airway regions at 19 pcw lungs. Ciliated cells (arrowhead), *FOXJ1*^+^/*SCGB3A2*^-^; secretory cells (arrow), *FOXJ1*^-^/*SCGB3A2*^+^. (I) MUC16^+^ ciliated cells (dashed line), ciliated cells (dashed circle), and secretory cells (arrow) located in the proximal cartilaginous airway regions of fetal lungs at 19 pcw. The MUC16^+^ ciliated cells express *MUC16* (white) with a weak level of *FOXJ1* (red), whereas the ciliated cells only express strong *FOXJ1* without *MUC16* expression. *SCGB3A2* (green) (J, J’) Proximal basal cells (J, dashed line) line the basal layer of the proximal cartilaginous pseudostratified airway in fetal lungs at 19 pcw and are marked by *TP63* (red), *F3* (white), and *IGFBP3* (green). In contrast, only a few *TP63*^+^ basal cells (J’, red, arrowheads) are observed in the non-cartilaginous, non-pseudostratified airway regions. (K) *ASCL1*^+^ pulmonary neuroendocrine (arrow) and MUC5AC^+^/ASCL1^+^ progenitors (arrowhead) in the non-cartilaginous airway regions of fetal lung at 12 pcw. MUC5AC (green), *ASCL1* (red), *SCGB3A2* (white). DAPI, nuclei. Scale bars, 50 μm. (L) Diagram describing spatial location of epithelial cell types observed in the developing human lungs.

**Figure S5.**
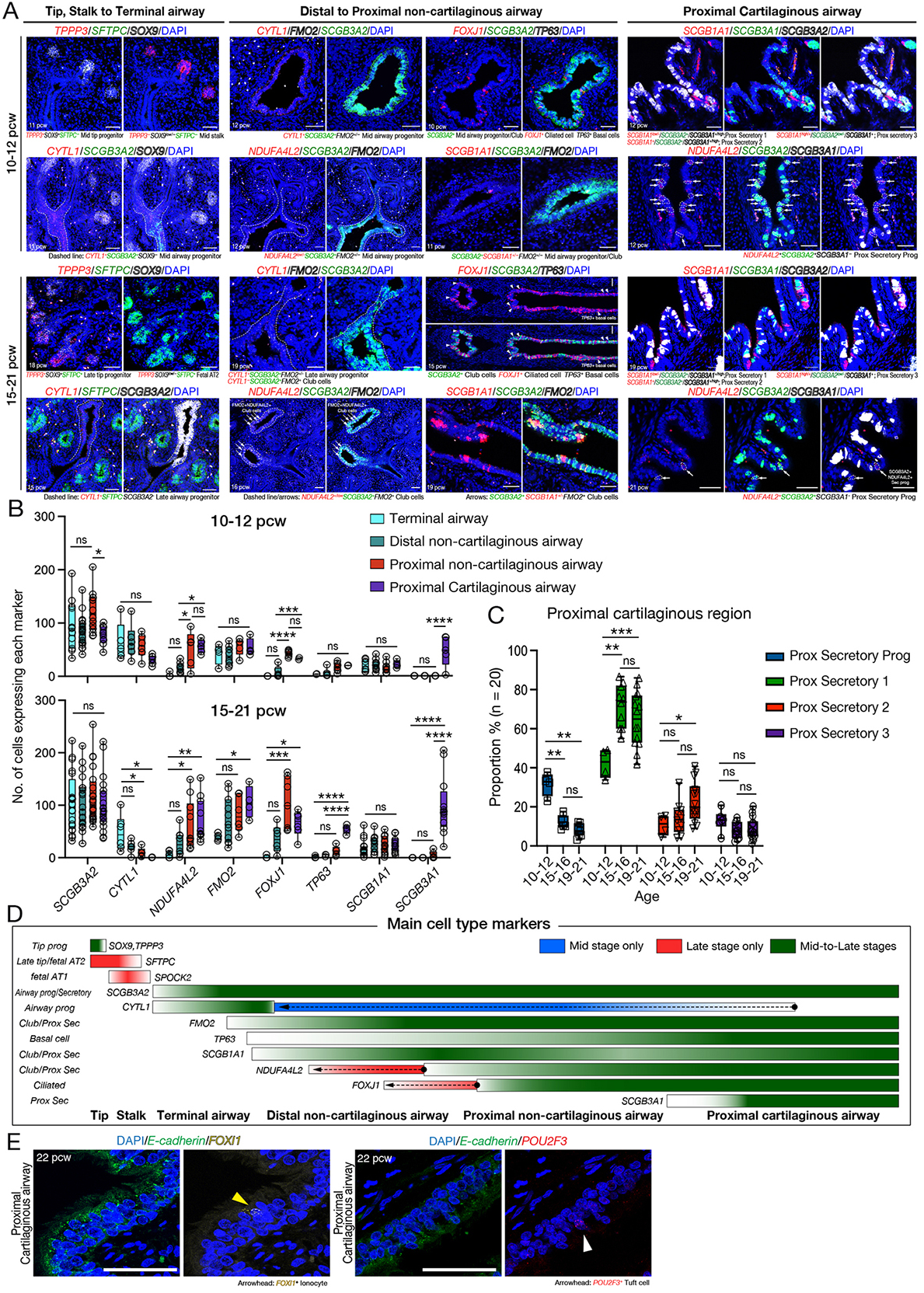
Spatiotemporal location, distribution, and quantification of major epithelial cell types along the distal to proximal axis of the developing lungs. (A) *In situ* HCR analysis of fetal human lung tissues at mid (10-12 pcw) and late (15-21 pcw) stages, showing spatiotemporal location and distribution of major epithelial cell types along the distal to proximal axis of the developing lungs. The lung regions were divided for imaging into tip, stalk to terminal airway; distal to proximal non-cartilaginous airway; and proximal cartilaginous airway. (B) Quantification of cells expressing marker genes of airway lineages along the airway regions at mid (10-12 pcw, upper) and late (15-21 pcw, lower) stages. *SCGB3A2*, airway progenitors/all secretory lineage cells; *CYTL1*, airway progenitor cells; *NDUFA4L2*, club/proximal secretory cells; *FMO2*, club/proximal secretory cells; *FOXJ1*, ciliated cells; *TP63*, basal cells; *SCGB1A1*, club/proximal secretory cells; *SCGB3A1*, proximal secretory cell subtypes 1-3. Significance was evaluated by 1-way ANOVA with Tukey multiple comparison post-test; n=3 biological replicates; ns: not significant \**P*<0.05, \*\**P*<0.01, \*\*\**P*<0.001, \*\*\*\**P*<0.0001. (C) Proportion of proximal secretory progenitor cells, proximal secretory cell subtypes 1-3 within the proximal cartilaginous airway regions by ages, at 10-12, 15-16, and 19-21 pcw. The secretory cells in the proximal cartilaginous airway regions were counted: Prox Secretory Prog, *SCGB3A2*^+^*SCGB3A1*^-^*SCGB1A1*^-^; Prox Secretory 1, *SCGB3A2*^+^*SCGB3A1*^+^*SCGB1A1*^-^; Prox Secretory 2, *SCGB3A2*^+^*SCGB3A1*^+^*SCGB1A1*^+^; Prox Secretory 3, *SCGB3A2*^-^ *SCGB3A1*^+^*SCGB1A1*^+^. Club cells located in the non-cartilaginous airway regions were excluded. Significance was evaluated by 2-way ANOVA with Tukey multiple comparison post-test; n=4 biological replicates; ns: not significant \**P*<0.05, \*\**P*<0.01, \*\*\**P*<0.001. (D) Diagram describing spatiotemporal distribution of major cell type markers along the distal to proximal axis of the developing lungs, at mid and late stages. Mid stage only, blue; Late stage only, red; Mid-to-late stages, green. Arrows indicate narrowed (*CYTL1*) or expanded (*NDUFA4L2*, *FOXJ1*) distribution after mid to late stage transition. (E) *In situ* HCR analysis of rare cell type markers of putative ionocytes (*FOXI1* yellow) and tuft cells (*POU2F3*, red). *E-cadherin*, green. DAPI, nuclei. Scale bar, 50 μm.

**Figure S6.**
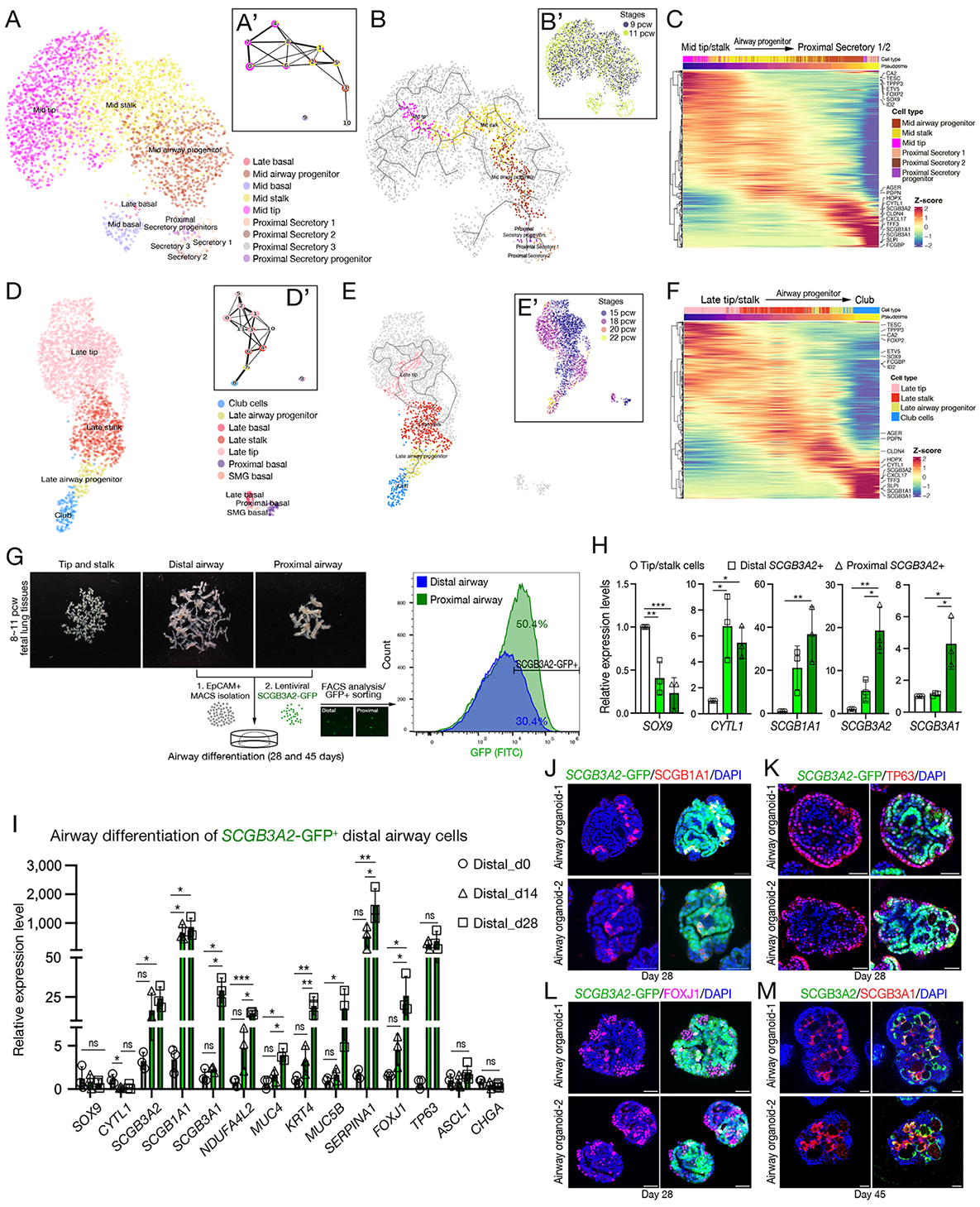
Trajectory analysis of airway lineage differentiation via airway progenitor cells in the developing human lung. (A, A’) UMAP visualization (A) and PAGA analysis (A’) of a lineage trajectory from mid tip to proximal secretory lineage cells, including proximal secretory progenitor and proximal secretory cell subtypes 1 to 3. Mid and late basal cells were shown to be disconnected from other proximal secretory cell types in the PAGA analysis (A’). (B-C) Trajectory UMAPs, by cell type (B) and stages (B’), and the relevant gene expression heatmap (C) displaying the selected lineage trajectory from mid tip to proximal secretory cell subtypes 1 and 2, analysed by Monocle 3. (Note that the grey lines in UMAP indicate all of the predicted differentiation paths from a user-defined starting point.) (D, D’) UMAP visualization (D) and PAGA analysis (D’) of a lineage trajectory from late tip, late stalk, late airway progenitor to club cells. Basal cells, including late basal, proximal basal, and SMG basal cells were shown to be left out of the trajectory as they do not connect clearly to the other cell types in this analysis (D’). (E-F) Trajectory UMAPs, by cell types (E) and stages (E’), and the relevant gene expression heatmap (C) showing the selected lineage trajectory from late tip to club cells, analysed by Monocle 3. (G) Purification of distal *SCGB3A2*-GFP^+^ airway cells from human fetal lung tissues at 8-11 pcw. The epithelial cells were isolated using EPCAM magnetic microbeads (MACS) from the dissected distal and proximal airway tissues, followed by infection with lentivirus habouring *SCGB3A2* promoter-driven GFP. The *SCGB3A2*-GFP positive cell fractions were sorted and analysed by FACS after 48 hrs and *in vitro* cultured for 28 and 45 days in the airway differentiation medium. (H) Gene expression profile of the freshly purified *SCGB3A2*-GFP positive cells derived from distal and proximal airway tissues were investigated by qRT-PCR and compared with dissected tip cells. *SOX9*, distal tip progenitor marker. *CYTL1*, airway progenitor marker. *SCGB1A1* and *SCGB3A2*, airway/secretory cell lineage markers. *SCGB3A1*, proximal secretory cell marker. Data was normalised to *SCGB3A2*-GFP negative cells derived from distal tip/stalk tissues; mean ± SD of 3 biological replicates. Significance was evaluated by 1-way ANOVA with Tukey multiple comparison post-test; \**P*<0.05, \*\**P*<0.01, \*\*\**P*<0.001. (I) Gene expression analysis of the *in vitro* cultured *SCGB3A2*-GFP positive cells (airway progenitors) derived from distal airway tissues by qRT-PCR. Airway organoids were formed from the *SCGB3A2*-GFP positive cells and collected at Day 0, 14, and 28 days after culture for the analysis. Data were normalised to *SCGB3A2*-GFP negative cells derived from distal tip/stalk tissues; mean ± SD of 4 biological replicates. Significance was evaluated by 1-way ANOVA with Tukey multiple comparison post-test; ns: not significant \**P*<0.05, \*\**P*<0.01.(J-M) Immunofluorescence analysis of two biologically independent, SCGB3A2-GFP+ cell-derived airway organoids cultured in the airway differentiation medium for 28 (J-L) and 45 (M) days. SCGB1A1 (J, red), airway progenitor/secretory cell marker. TP63 (K, red), basal cell marker. FOXJ1 (L, magenta), ciliated cell marker. SCGB3A1 (M, red), proximal secretory cell marker. DAPI, nuclei. Scale bar, 50 μm.

**Figure S7.**
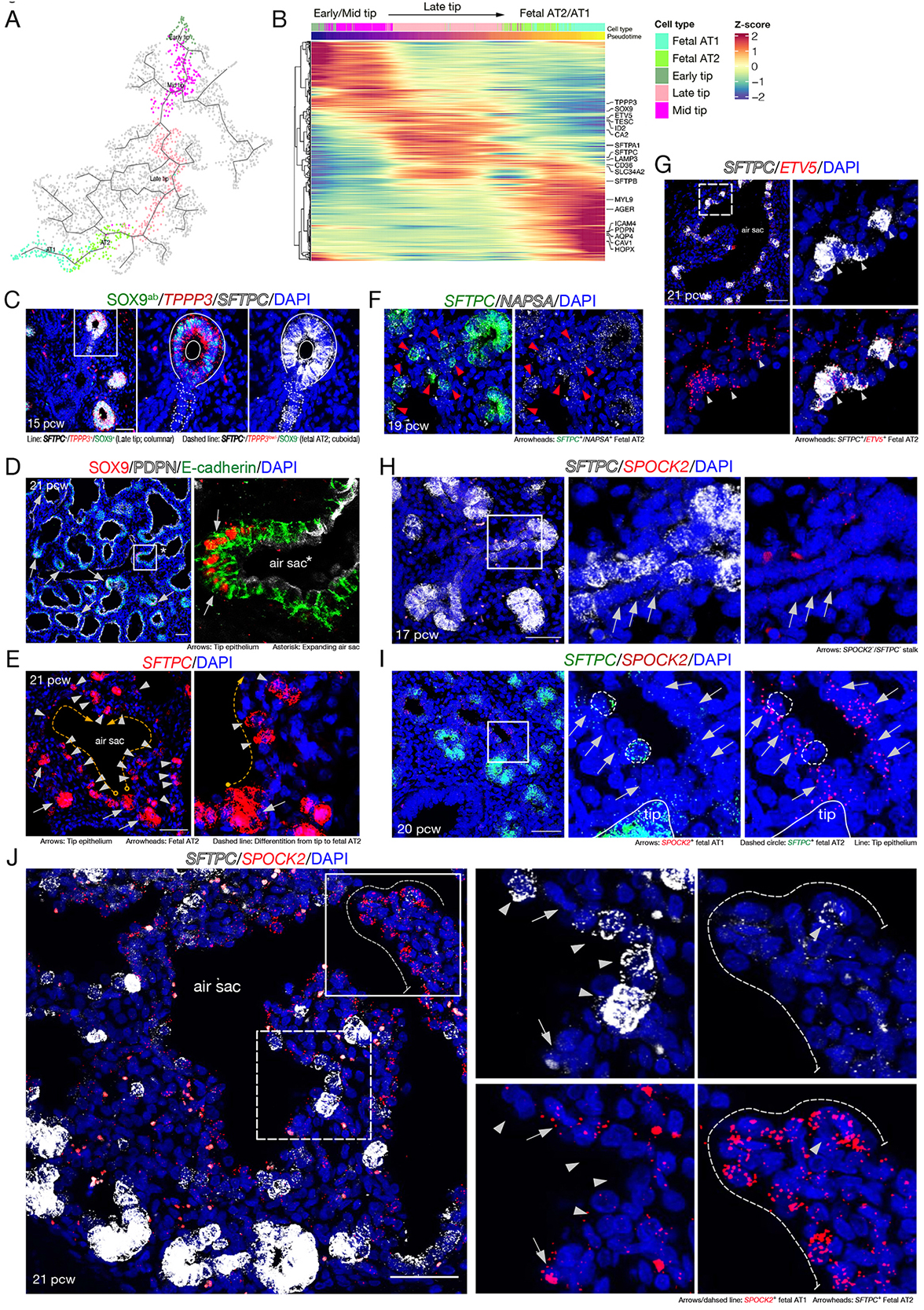
Late epithelial tip cells differentiate to AT2 and AT1 cells. (A, B) UMAP visualisation (A) of a lineage trajectory from early/mid/late tip to fetal AT2 and AT1 cells and the relevant gene expression heatmap (B) showing the selected lineage trajectory analysed by Monocle 3. (C) *In situ* HCR (*TPPP3* and *SFTPC*) and immunostaining (SOX9) analysis of 15 pcw fetal lung, describing SOX9^+^*TPPP3*^+^*SFTPC*^+^ tip epithelial progenitors (lines) and SOX9^-^*TPPP3*^-^*SFTPC*^+^ fetal AT2 cell population (dashed circles) lining the stalk. (D) Immunostaining of 21 pcw fetal lung using antibodies against SOX9 (red), PDPN (white), and E-cadherin (green). Arrows indicate the late tip cell population, which does not co-express the stalk marker, PDPN. (E-G) *In situ* HCR analysis of 19 (F) and 21 pcw (E, G) fetal lungs, showing the *SFTPC*^+^ fetal AT2 cell population (arrowheads) lining the developing air sacs. Arrows indicate *SFTPC*^+^ late tip cells. (E) *SFTPC* (red). (F, G) *NAPSA* (white; F) and *ETV5* (red; G) overlap with *SFTPC* in the fetal AT2 cells. (H-J) *In situ* HCR analysis of distal lung regions at 17 (H), 20 (I), and 21 (J) pcw, visualising *SFTPC*^-^ fetal AT1 cells (arrows). *SFTPC*^-^*/SPOCK2*^-^ stalk cells at 17 pcw (H) began to express *SPOCK2* (red) at 20 pcw (I) and further developed to future AT1 cells (*SFTPC*^-^*/SPOCK2*^+^) at 21 pcw (J). Dashed circles (I) and arrowheads (J) indicate fetal AT2 cells. Dashed line (J) shows fetal AT1 cells lining the developing air sacs. DAPI, nuclei. Scale bars, 50 μm.

**Figure S8.**
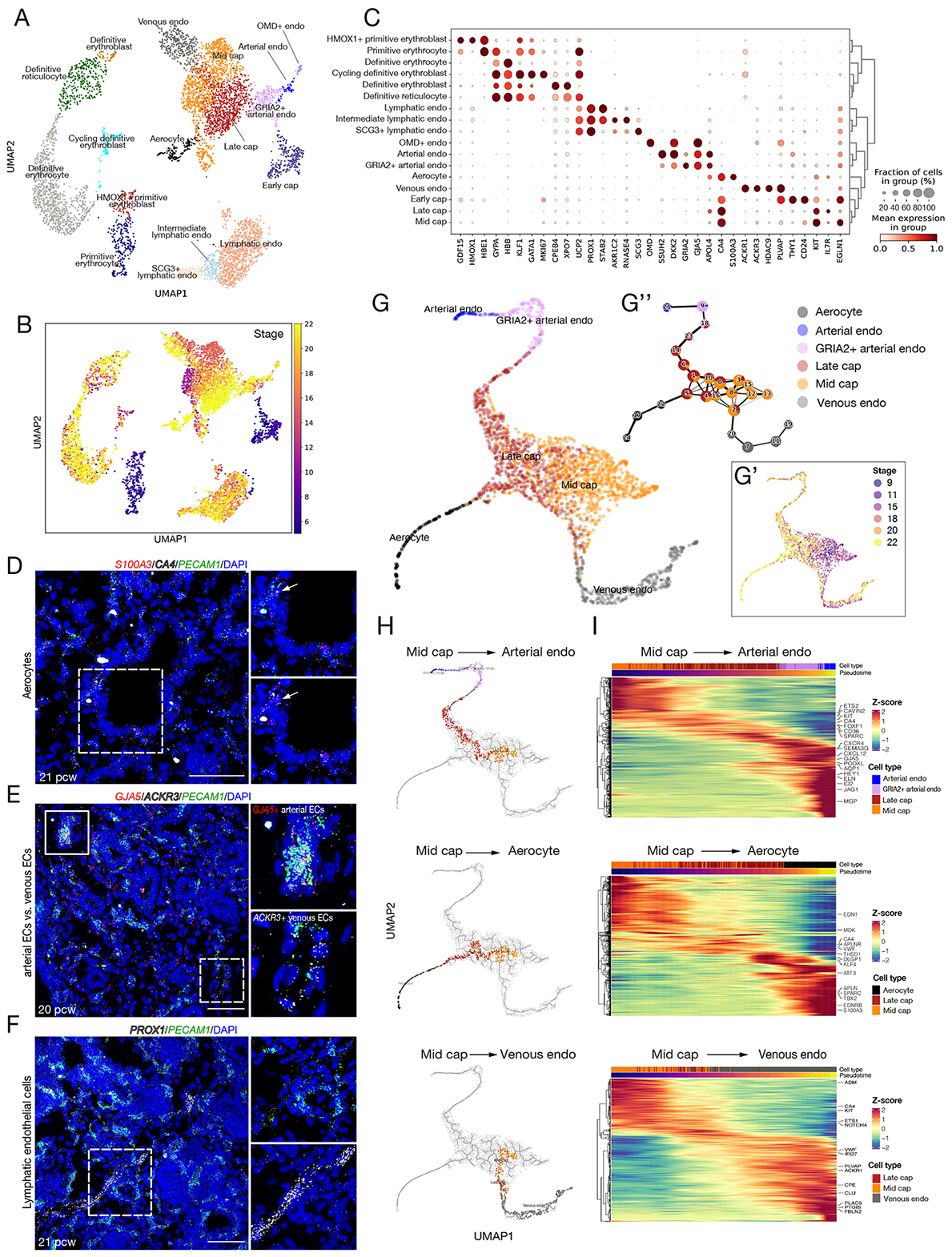
Endothelial cell types in the developing human lung. (A,B) UMAP visualisation of endothelial cells, coloured by cell types (A) and stages (B). (C) Dot plot describing differential marker gene expression level by cell type. (D-F) *In situ* HCR analysis of distal lung regions at 20 (E), and 21 (D, F) pcw. (D) Aerocytes (*S100A3*^+^ red/*CA4*^+^ white), capillary endothelium (*CA4*^+^ white), and all endothelial cells (*PECAM^+^*, green). (E) Arterial endothelial cells (*GJA5*^+^ red), venous endothelial cells (*ACKR3*^+^ white), and all endothelial cells (*PECAM^+^*, green). (F) Lymphatic endothelial cells (*PROX1*^+^ white) and all endothelial cells (*PECAM^+^*, green). DAPI, nuclei. Scale bars, 50 μm. (G) Trajectory UMAP and PAGA plot (G’’) visualising potential endothelial cell lineage hierarchy from Mid/Late capillary endothelial cells to arterial endothelial cells, aerocytes, or venous endothelial cells coloured by cell types (G) and stages (G’). (H, I) Individual trajectory UMAPs (H) and the relevant gene expression heatmaps (I) displaying potential lineage trajectories derived by Monocle 3 from Mid/Late capillary endothelial cells to arterial endothelial cells (*top*), aerocytes (*middle*), or venous endothelial cells (*bottom*).

**Figure S9.**
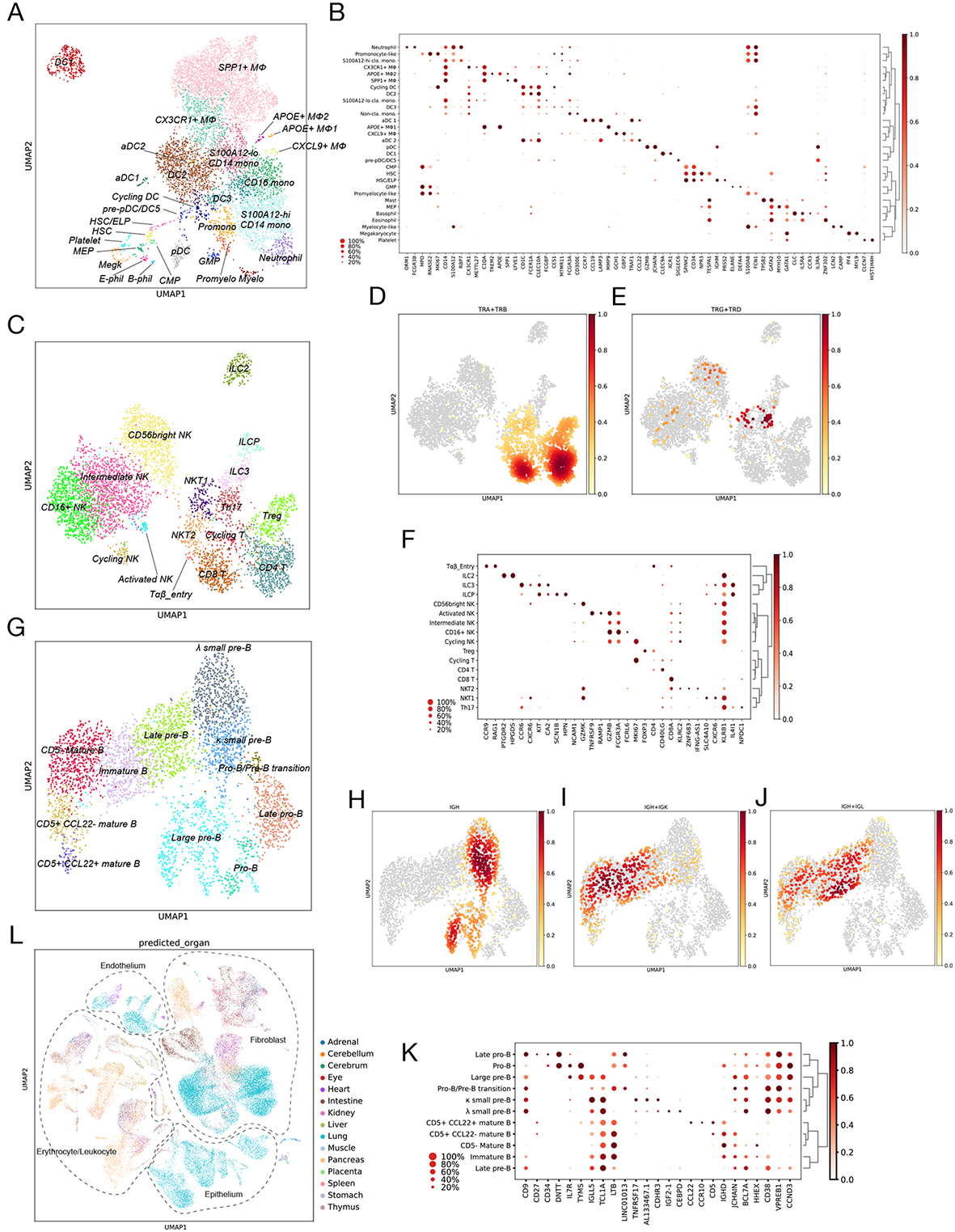
Clustering and cell type markers for immune cell types and comparison to other fetal data sets. (A,C,G) UMAP embeddings of different immune compartments showing myeloid cell types/states (A), T, NK and ILC lymphoid cells (C) and B lymphoid cells (G). (B,F,K) Dot plots showing expression of selected marker genes of cell types/states in the three immune compartments. (D, E, H, I, J) Enrichment of each class of immune receptors based on abTCR, gdTCR and BCR-enriched scRNA-seq. (L) Predicted organ-of-source with highest scores for cells shown in Figure 1, based on the reference atlas in (Cao et al. 2020).

**Figure S10.**
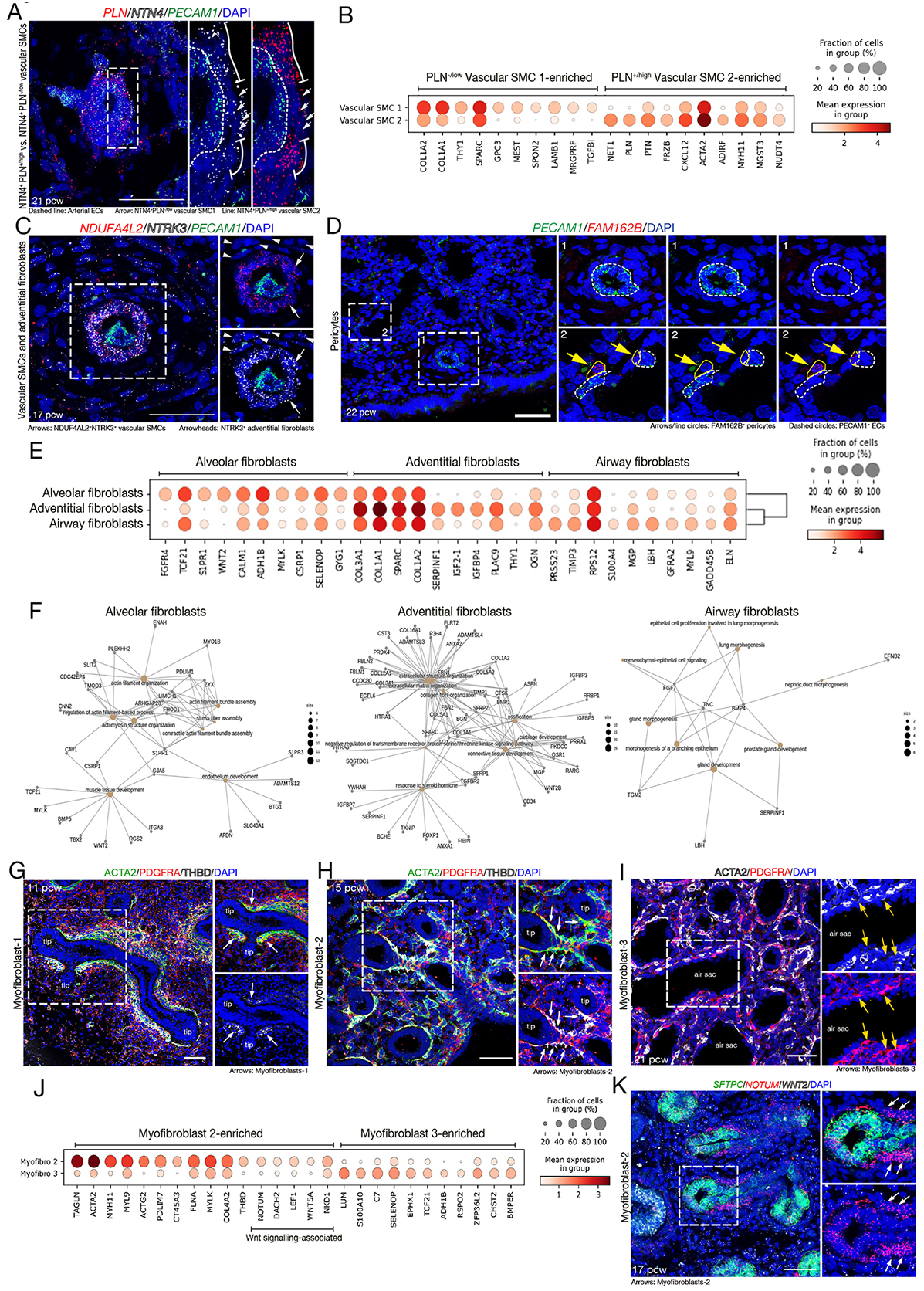
Spatial analysis of mesenchymal cell types in the developing human lungs by *in situ* HCR assay and immunostaining. (A) Vascular SMC 1 and 2 are surrounding arterial endothelial cells (*PECAM1*^+^, dashed line), each marked by *NTN4*^+^/*PLN*^-/low^ (vSMC 1, arrows) and *NTN4*^+^/*PLN*^+/high^ (vSMC 2, lines). (B) Dot plot describing differential gene expression between vascular SMC 1 and 2. (C) Vascular SMCs and adventitial fibroblasts in 17 pcw fetal lung. *NDUF4AL2*^+^ red/*NTRK3*^+^ vSMCs (arrows) are surrounded by *NDUF4AL2*^-^/*NTRK3*^+^ adventitial fibroblasts (arrowheads). *PECAM1* (green) indicates an endothelial cell tube. (D) *FAM162B*^+^ pericytes (red) are surrounding *PECAM1*^+^ endothelial cells (green) in the microvascular regions. (E) Dot plot describing differential marker gene expression level between alveolar, adventitial and airway fibroblasts. (F) Concept network visualisation of gene ontology (GO) analysis using clusterProfiler for differentially expressed genes in alveolar, adventitial and airway fibroblasts. (G-I) Immunostaining of fetal lung tissues at 11 (G), 15 (H), and 21 (I) pcw, to visualise myofibroblast populations: Myofibroblast-1 (G) and -2 (H) surrounding the developing stalk epithelial tubes, and Myofibroblast-3 (I) surrounding the developing air sacs. ACTA2^+^/PDGFRA^+^ Myofibroblast-1 (THBD^weak^; G) and -2 (THBD^high^, arrows; H). PDGFRA^+^ Myofibroblast-3 at 21 pcw, does not express ACTA2 (arrows; I). (J) Dot plot describing differential gene expression level between myofibroblast-2 and -3. The myofibroblast-2 population showed enriched expression of Wnt signalling associated genes, e.g. *NOTUM*, *LEF1*, and *DACH2*. (K) *In situ* HCR assay of 17 pcw fetal lung tissues. Myofibroblast-2 expresses *NOTUM* (red), a Wnt antagonist, to block local Wnt signals from alveolar fibroblasts (white, *WNT2*) to the stalk epithelium. DAPI, nuclei. Scale bars, 50 μm.

**Figure S11.**
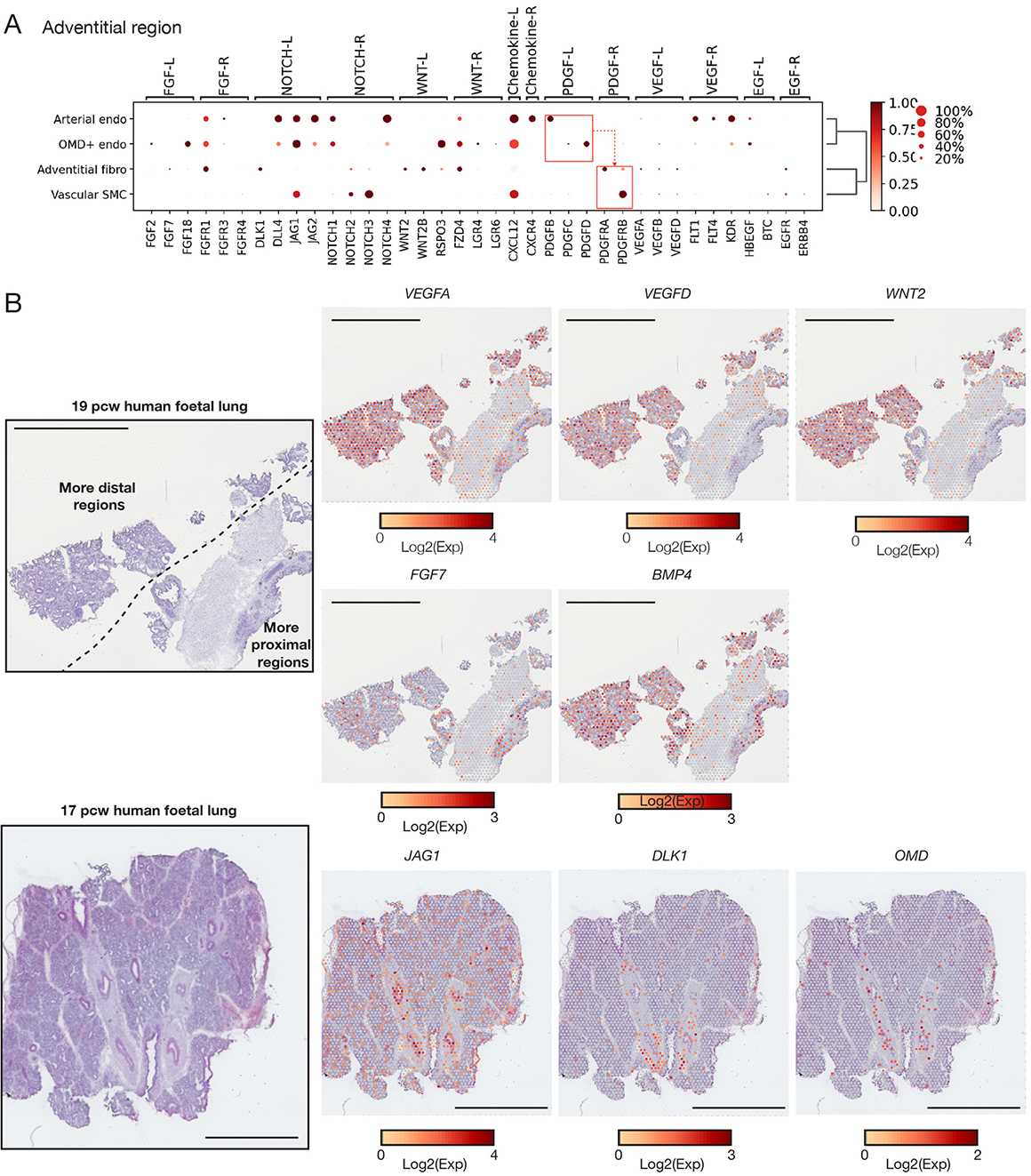
Signalling ligand-receptor interactions in the adventitial niche. (A) Overview of predicted ligand–receptor interactions using CellPhoneDB in the adventitial niche. Dot plots visualise gene expression by cell type and dashed arrows indicate a predicted direction of signalling from ligands to receptors. (B) Visium spot transcriptome cluster map visualising signalling ligands expressed in the fetal lung tissues at 19 (upper) and 17 (lower) pcw. Scale bars denote 2.5 mm (upper) and 2 mm (lower), respectively.

**Figure S12.**
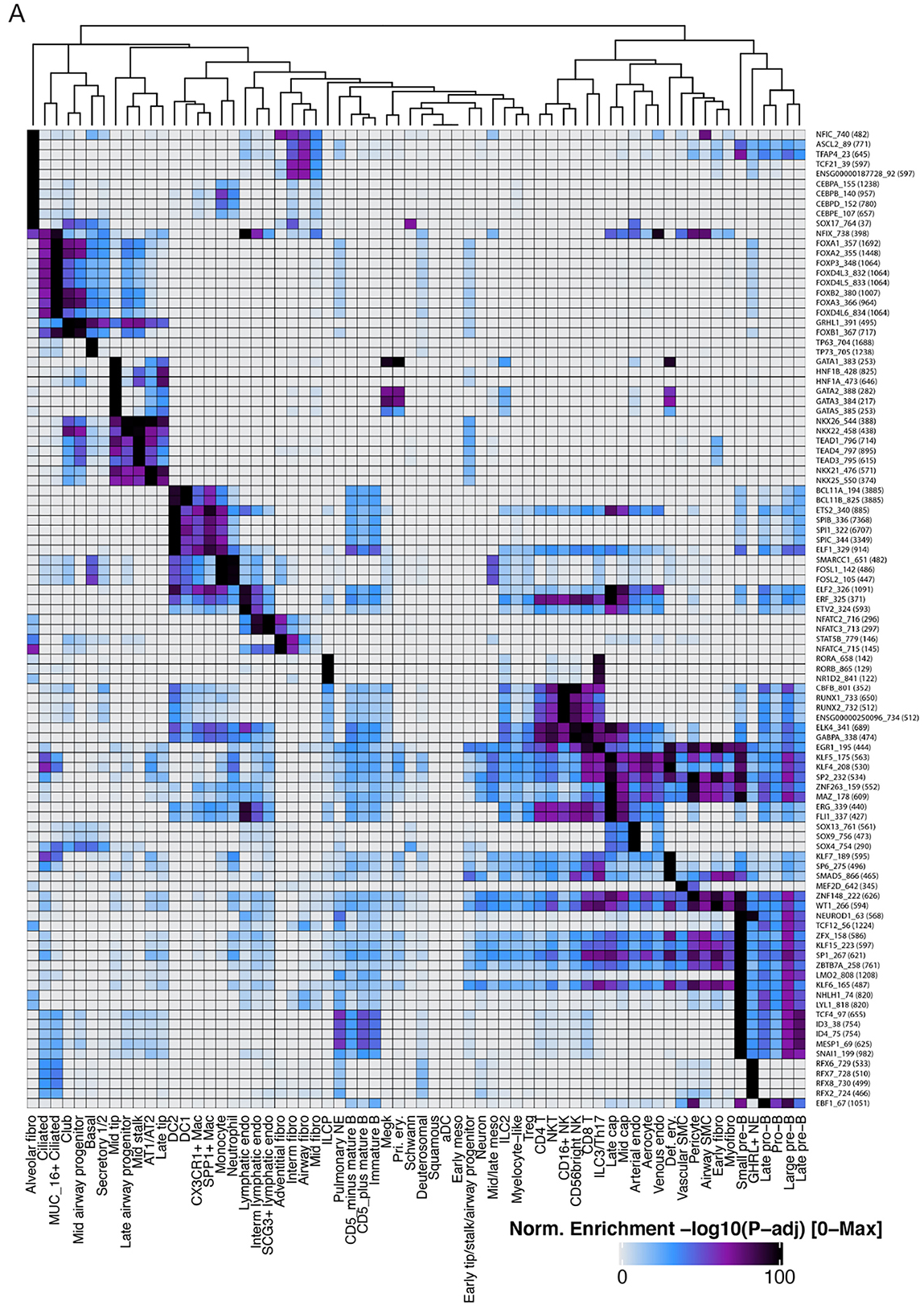
Global landscape of motif enrichment. Top 5 enriched motifs in the marker peaks among all the cell types/states. Statistical significance is visualised as a heatmap according to the colour bar below.

**Figure S13.**
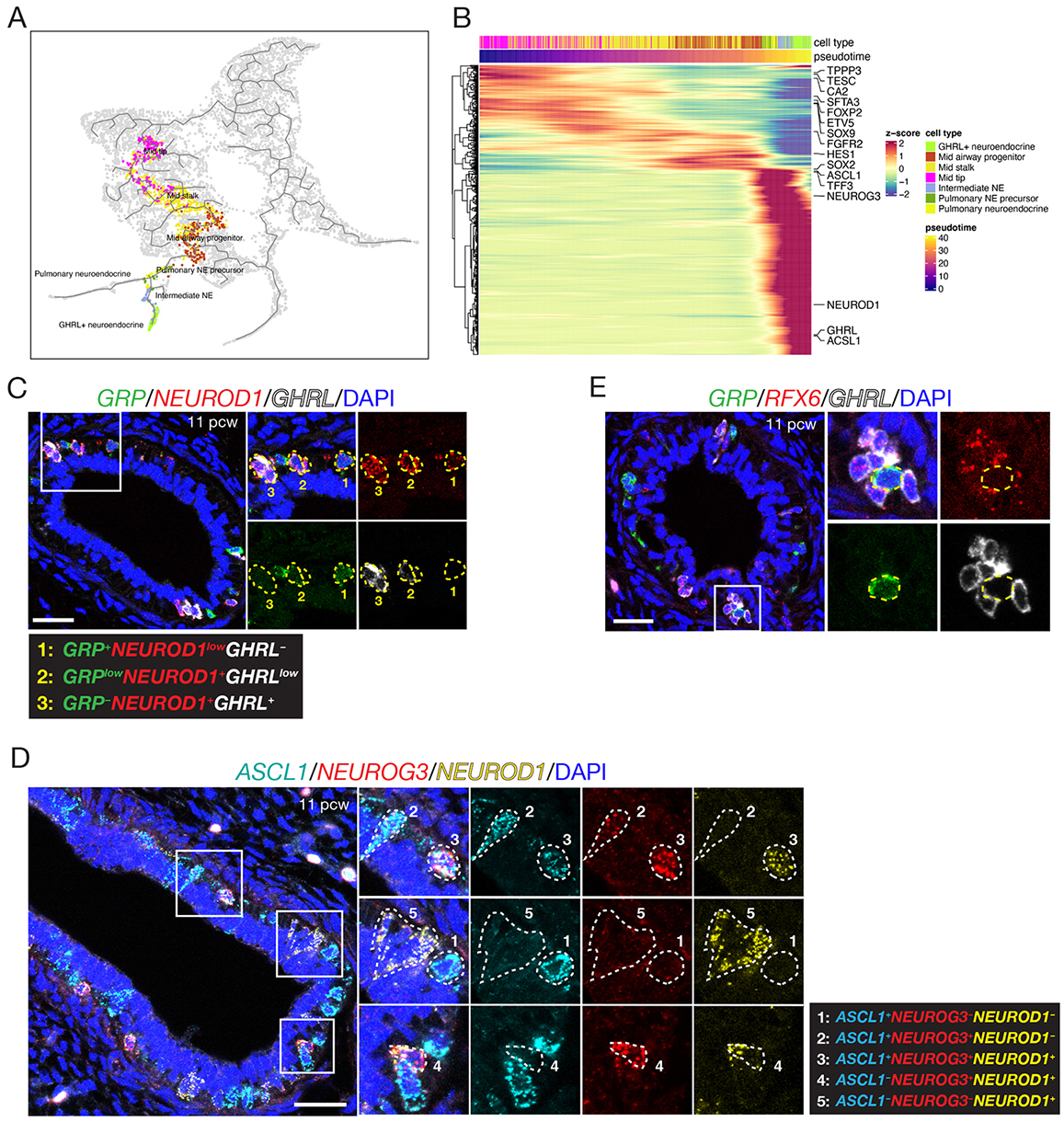
Transcription factor regulatory network controlling NE subtypes. (A) Selected trajectory from Mid tip cells to *GHRL^+^* NE cells via Intermediate NEs, a transition cell population. (B) Heatmap of genes differentially expressed along the trajectory. (C) Representative HCR images showing the transition between two types of NE cells. *GRP* (green), *NEUROD1* (red), *GHRL* (white). #1 labelled *GRP^+^NEUROD1^low^GHRL^-^* cells, which have just started the transition from GRP^+^ pulmonary NE/precursor cells. #2 labelled *GRP^low^NEUROD1^+^GHRL^low^* cells, in transition to *GHRL^+^* NE cells. #3 labelled *GRP^-^ NEUROD1^+^GHRL^+^*, *GHRL^+^* NE cells. Right: Mean ± SEM of *NEUROD1^+^*cell types. 11 pcw: N=2 fetal lungs, n = 129 *NEUROD1^+^* cells; 12 pcw N=3 fetal lungs, n=132 *NEUROD1^+^* cells. Scale bars = 25 μm in all panels. (D) Representative HCR images showing *NEUROG3* co-expression with *ASCL1* and *NEUROD1*. Dashed white lines label representative cells showing different combinations of the three transcription factors, further indicated by #1-#5 labelling. *ASCL1* (cyan), *NEUROG3* (red), *NEUROD1* (yellow). (E) Representative HCR images showing *RFX6* expression in *GHRL^+^* NE cells. Dash yellow line labelled *GRP^+^RFX6^-^* pulmonary NE cells. Scale bars = 25 μm in all panels.

**Figure S14.**
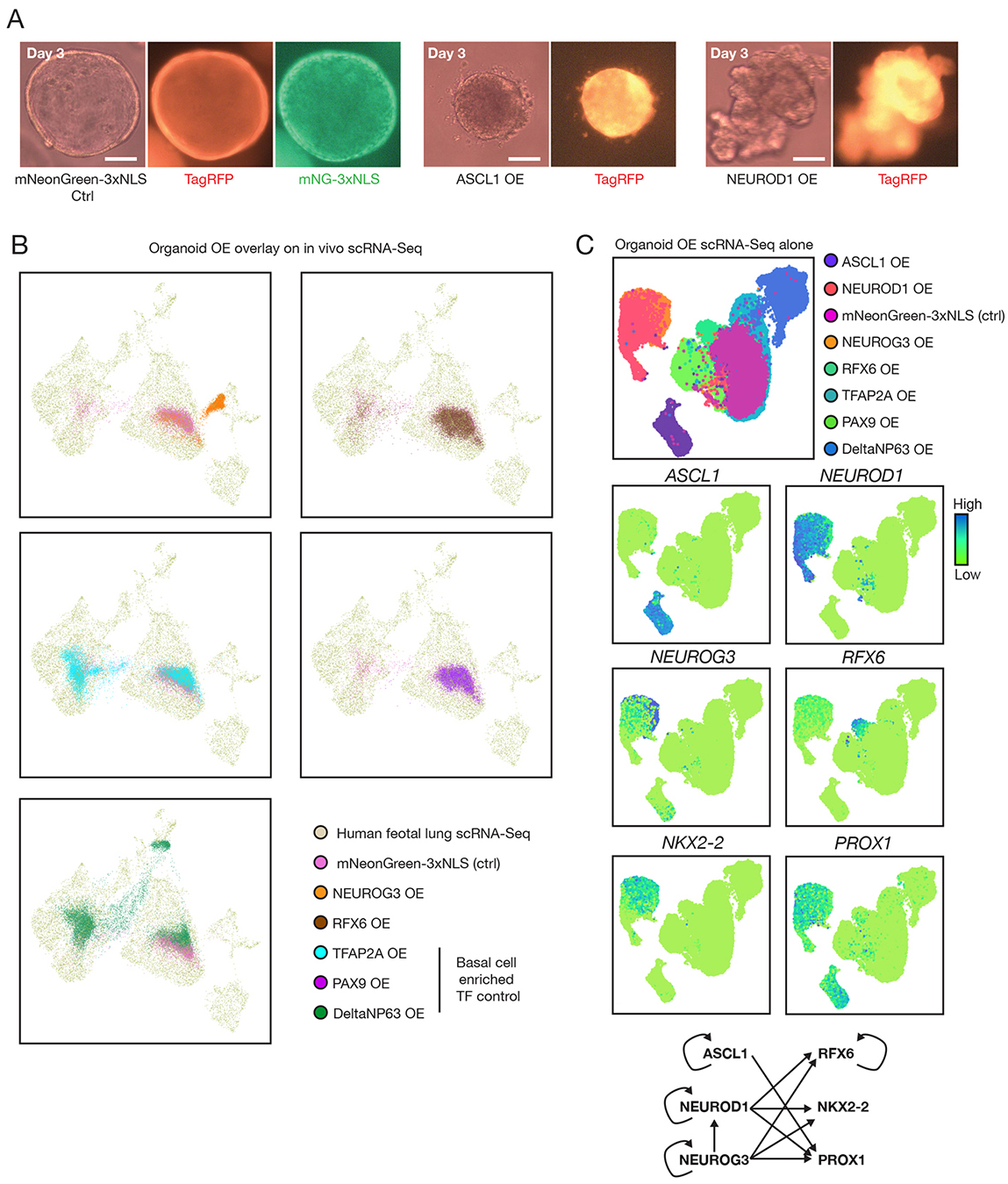
Validation of NE transcription factors using human fetal lung organoid system. (A) Representative epifluorescent microscopic images showing organoid morphology after 3 days of mNeonGreen-3xNLS (control), ASCL1, or NEUROD1 overexpression. (B) ScRNA-seq results of organoid transcription factor overexpression overlay on human fetal lung scRNA-seq as a reference. (C) scRNA-seq results of transcription factor overexpression; organoid data only in the UMAP. Selected transcription factor expression was shown in the middle panel. A regulatory network of the selected transcription factors were drawn based on the organoid OE data at the bottom of the panel. (Note that the arrows do not necessarily denote direct interactions).

